# The genome of the Wollemi pine, a critically endangered “living fossil” unchanged since the Cretaceous, reveals extensive ancient transposon activity

**DOI:** 10.1101/2023.08.24.554647

**Authors:** Dennis Wm. Stevenson, Srividya Ramakrishnan, Cristiane de Santis Alves, Laís Araujo Coelho, Melissa Kramer, Sara Goodwin, Olivia Mendevil Ramos, Gil Eshel, Veronica M. Sondervan, Samantha Frangos, Cecilia Zumajo-Cardona, Katherine Jenike, Shujun Ou, Xiaojin Wang, Yin Peng Lee, Stella Loke, Maurizio Rossetto, Hannah McPherson, Sebastiano Nigris, Silvia Moschin, Damon P. Little, Manpreet S. Katari, Kranthi Varala, Sergios-Orestis Kolokotronis, Barbara Ambrose, Larry J. Croft, Gloria M. Coruzzi, Michael Schatz, W. Richard McCombie, Robert A. Martienssen

**Author notes:** These authors contributed equally (Dennis Wm. Stevenson, Srividya Ramakrishnan, Cristiane de Santis Alves, Laís Araujo Coelho, Melissa Kramer).

## Abstract

We present the genome of the living fossil, *Wollemia nobilis*, a southern hemisphere conifer morphologically unchanged since the Cretaceous. Presumed extinct until rediscovery in 1994, the Wollemi pine is critically endangered with less than 60 wild adults threatened by intensifying bushfires in the Blue Mountains of Australia. The 12 Gb genome is among the most contiguous large plant genomes assembled, with extremely low heterozygosity and unusual abundance of DNA transposons. Reduced representation and genome re-sequencing of individuals confirms a relictual population since the last major glacial/drying period in Australia, 120 ky BP. Small RNA and methylome sequencing reveal conservation of ancient silencing mechanisms despite the presence of thousands of active and abundant transposons, including some transferred horizontally to conifers from arthropods in the Jurassic. A retrotransposon burst 8-6 my BP coincided with population decline, possibly as an adaptation enhancing epigenetic diversity. *Wollemia*, like other conifers, is susceptible to *Phytophthora*, and a suite of defense genes, similar to those in loblolly pine, are targeted for silencing by sRNAs in leaves. The genome provides insight into the earliest seed plants, while enabling conservation efforts.

## Introduction

The discovery of the Wollemi pine (*Wollemia nobilis*) and its description by Jones et al. ^1^ was to the studies of gymnosperm biology analogous to that of the Coelacanth to Ichthyology in 1939. It was recognized immediately as being something unusual that until then had been known only from the fossil record ^2,3^. This resulted in Wollemi pine becoming designated a “living fossil” - a term originally coined by Darwin in “On The Origin of Species”. Known from only four nearby localities in steep nearly inaccessible canyons, the wild population of *Wollemia nobilis* consists of approximately 60 individual trees and associated ramets, tightly confined to a small region of Wollemi National Park approximately 100 km west of Sydney, Australia, and were threatened by the unprecedented bushfires of 2019. An initial population genetic survey could find no genetic differences between or within any of the populations ^4^. *Wollemia* cuttings were found to be highly susceptible to phytophthora infection, which is shared with other conifers and common to many Australian native plants. *Wollemia* was classified as critically endangered (CR) on the IUCN’s Red List in 1998 ^5^ and conservation has been a priority since ^6^.

*Wollemia* is well known in the southern hemisphere fossil record with leaves, twigs or pollen found in South America ^7^, Antarctica ^8^, New Zealand ^9^ and Australia ^10^, ^11^ from approximately 90 to 2.5 My before present (BP). The large range and frequency of fossils and pollen are strong evidence for *Wollemia* being abundant even up to several million years BP. Pollen morphology of *Wollemia* matches the fossil pollen morphogenus *Dilwynites* ^12^, though it is unclear to what degree the *Dilwynites* group also comprises the morphologically close *Agathis* based on palynological evidence. *Dilwynites* pollen is found in Antarctica, South America, and Bass strait sea bed cores ^7,13,14^, which confirms the frequency of fossil finds. *Agathoxylon*, which shares many *Wollemia* traits, is found from the late paleozoic 250 My BP^15^ with a worldwide distribution, is the likely progenitor genus of *Wollemia*, and one of the earliest trees (Figure1).

**Figure 1.**
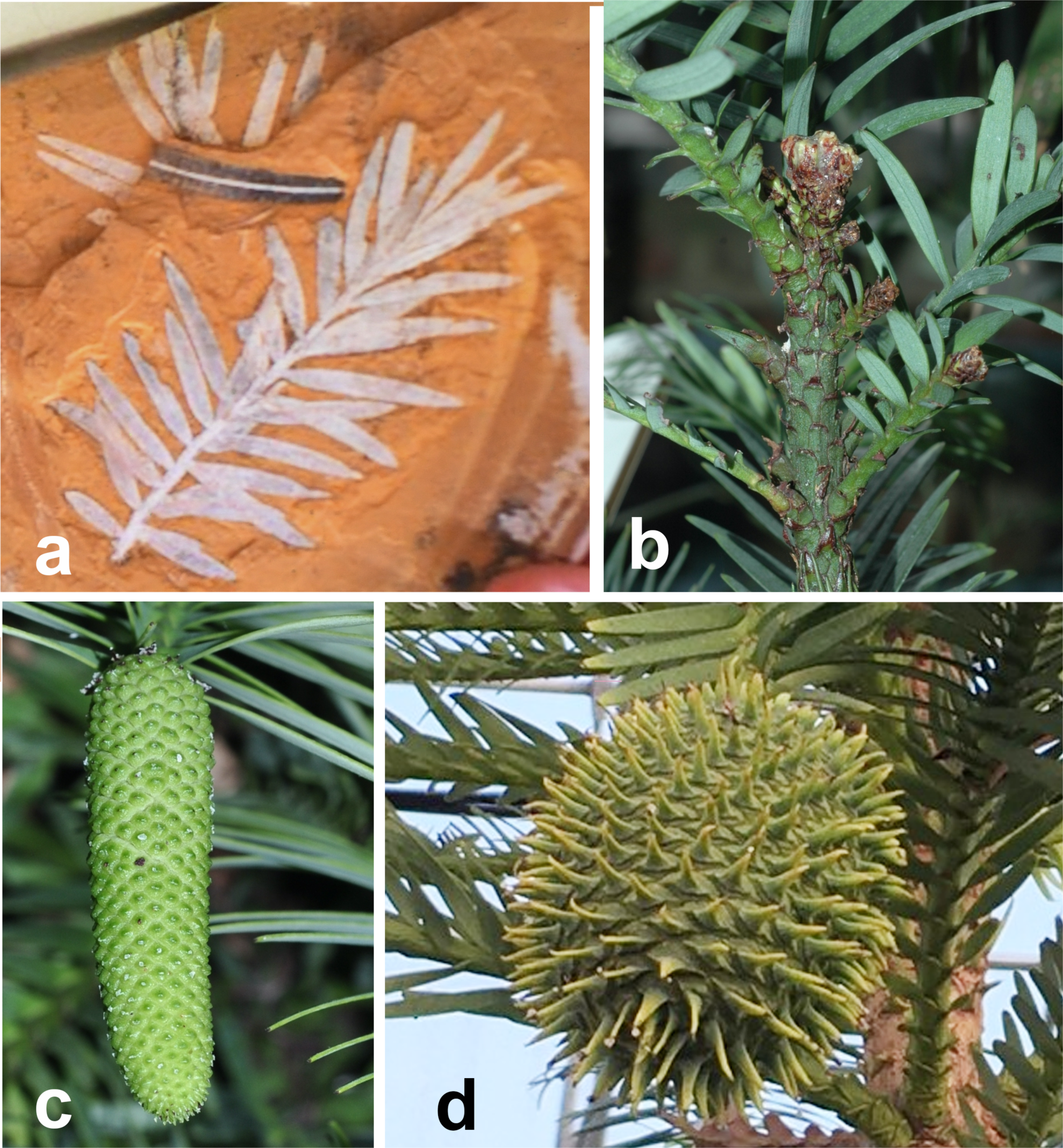
*Podozamites notabilis* and *Wollemia nobilis*. **a,** Shoot of *P. notabilis* a 150 million year old fossil relative of *Wollemia*. **b-d,** *Wollemia nobilis*. **b,** Vegetative shoot. **c,** Pollen strobilus. **d,** Ovulate strobilus.

The Wollemi pine is monoecious, with pollen cones developing from five years of age, ovulate cones from eight years of age, and the short-lived seeds are generally poorly dispersed ^16^. It has a prolific coppicing habit, even in absence of disturbance, with some individuals comprising over 40 trunks ^1^. Individual trunks can live in excess of 400-450 years ^17^ and may be replaced by other ramets after senescence. Previous genetic studies could only identify very low levels of nuclear and plastid genetic diversity ^18,19^, suggesting either a very recent bottleneck from an already restricted genetic pool, or long-term persistence as one or very few closely related clones. As part of the overall management and conservation of the Wollemi pine, a global meta collection and a translocation strategy informed by genetic data are currently being planned ^16^.

Since its discovery, there has been considerable interest and study on many aspects of *Wollemia* biology ranging from its phylogenetic position to anatomy, karyology, phytochemistry, growth habit, and potential for bioprospecting. Based on morphology, *Wollemia* was clearly recognized as a third member of the conifer family, Araucariaceae, along with *Araucaria* and *Agathis*. Following earlier studies, the largest most comprehensive evolutionary analysis used morphological and anatomical data from extant and fossil taxa combined with molecular data representing 23 genomic regions (19 plastid, two nuclear, and two mitochondrial), and found *Wollemia* and *Agathis* to form a clade sister to *Araucaria* ^20^. Furthermore, they were able to place fossil genera, *Wairarapaia mildenhallii* and *Emwadea microcarpa,* on the stem lineage of the *Wollemia* + *Agathis* clade. The close relationship of these species with *Agathis* and *Wollemia* was previously suggested ^21,12,22^, based on several shared features in the ovulate cones. Given fossil phylogenetic calibrations, the origin of Araucariaceae was estimated at 308 ± 53 My BP and the origin of the *Wollemia* + *Agathis* clade at 61 ± 15 My BP ^23^.

Of particular interest has been the architecture of the Wollemi pine because it has a unique mode of rhythmic rather than continuous growth – a previously unknown trait among Araucariaceae ^24–27^. Trees of Wollemi pine develop multiple coppice shoots and orthotropic epicormic shoots that can become replacement or additional leaders. Unlike most conifers, where the leaf axils generally do not have axillary buds, the Araucariaceae do have axillary buds, but those are not well differentiated from surrounding tissue ^28,29^. In Wollemi pine leaf axils, a discernible shell zone is found around a small group of slightly differentiated cells that is typical of axillary meristems ^28^. These axillary buds remain hidden because in origin they become buried beneath the bark by localized cork cambia. They display a slow continuous development and remain as a long-lived source for orthotropic epicormic shoots that may become replacement or additional leaders shoots as well as coppicing ^29^. The genetic basis for this remains unknown. There is a paucity of information on other aspects of *Wollemia* structure and development. Wood anatomy is consistent with Araucariaceae and no unique features are present in *Wollemia ^30^*.

The *Wollemia* chromosome number is 2*n*=26 with a karyotype composed of nine pairs of metacentric, two pairs of meta-submetacentric, and two sub telocentric chromosomes with a C-value of 13.94 pg ^31^. Some aspects of seed germination and seedling survival have been described ^32,33^. Field observations and laboratory experiments indicate that, following seed shed in summer and early autumn when temperatures are high, *Wollemia* seeds germinate, especially if exposed to light. The ungerminated seeds that are shed late in the season survive over winter and germinate once temperatures rise in the spring. Experimental data indicate that seed germination is enhanced with stratification corresponding to lower temperatures in the winter. Current data indicates that the adult population structure of Wollemi Pine is composed of a small number of genets reproducing vegetatively with new genets being added to the adult population from the young, understory seedlings when gaps in the canopy occur because of fire or death of a large parent tree.

Whereas the salient features of *Wollemia* anatomy, morphology, karyology etc. are well described, genetic aspects of growth, development, and reproduction are lacking. Thus, in order to further our understanding for the survival of Wollemi pine in a changing environment, a genomic approach is necessary and, in this case, begins with a whole genome sequence. In this way, we can begin to understand how this living fossil species has managed to survive the vicissitudes of time and utilize that for current climate challenges to its persistence and to that of other endangered species.

## Results

### *Wollemia nobilis* genome sequencing and assembly

Given the extreme genome size of *W. nobilis*, we adopted a long read sequencing strategy based on Oxford Nanopore Technologies (ONT). Genomic DNA was isolated from a clonal propagule of Wollemi pine at the New York Botanical Garden. Protocol optimizations allowed us to employ aggressive size selection to generate the very long fragments necessary for assembly contiguity and repeat resolution in this large 12 Gb genome (see **Methods**). Using this approach, the ONT reads had a read length N50 of ∼25 kb, and we produced 36× coverage of the genome in reads ≳30 kb (Supplementary Figure1). Furthermore, re-basecalling of the raw ONT data using the improved high accuracy model in Guppy v4 provided increased read quality to improve the accuracy of the assembled consensus quality (Supplementary Figure1). In addition, genome size and heterozygosity were estimated from the 21-mer counts of ∼20× short Illumina reads using KMC and Genomescope 2.0 ^34^ (Supplementary Figure1). This method yielded a genome size estimate of 12.2 Gb, which is consistent with the previous estimate of 13.88 Gb (2C = 28.4 pg) ^35^, as well as an exceptionally low level of heterozygosity (∼0.001%).

The initial contig assembly of *W. nobilis* using the Flye assembler ^36^ yielded a total of 11.66 Gbp in 23,120 contigs. This assembly was then polished using Medaka (https://github.com/nanoporetech/medaka) (2 rounds) using long Nanopore reads, 1 round of POLCA ^37^ using Illumina short reads and 2 rounds of Illumina kmer based polishing using our novel method HetTrek (see **Methods**). Further evaluation of the assembly using Merqury ^38^ with short Illumina kmers revealed a consensus quality score (QV) of 31 with ∼7 errors per 10 kb. This resulted in the highly contiguous genome assembly (*Wollemia nobilis* v1.0) with a contig N50 of 8.83 Mb (Supplementary Table S1.1). This represents one of the highest contig N50 of all gymnosperm genomes assembled to date (Figure 2a). The contiguity of *Wollemia nobilis* 1.0 is most apparent when comparing with other gymnosperm assemblies (Supplementary Table S1.2). *Wollemia nobilis* 1.0 has a contig N50 size of 8.83 Mb, which is over three times larger than the largest contig N50 reported earlier in *Pinus tabuliformis ^39^* and Taxus chinensis ^40^ and nearly an order of magnitude larger than other published conifers.

**Figure 2.**
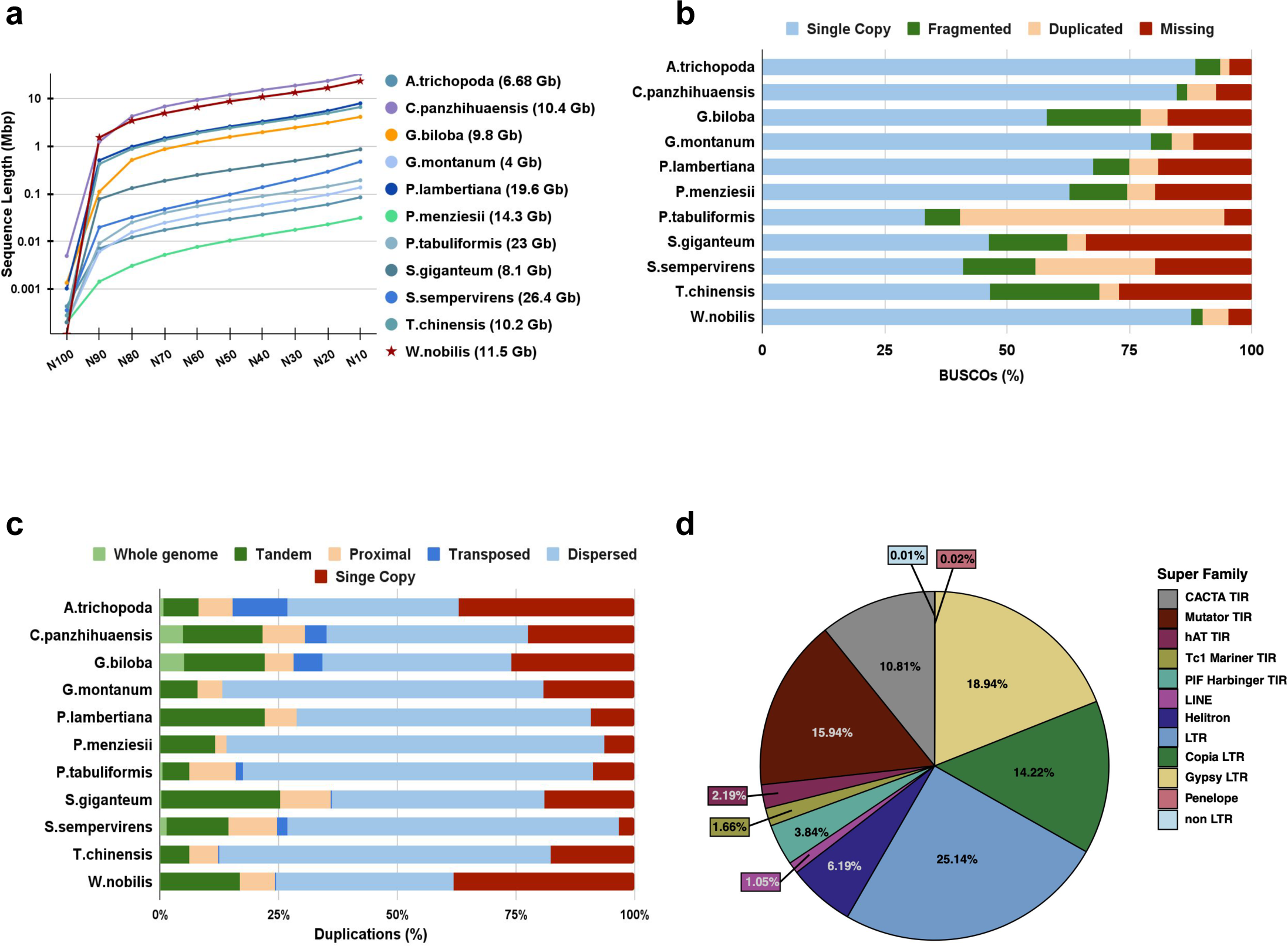
Genome Assembly and Annotation. **a,** Assembly contiguity of the *W. nobilis*1.0 genome, other gymnosperm genomes and *A. trichopoda* genome at the contig scale. **b,** BUSCO (Benchmarking Universal Single Copy Ortholog) gene set assessment of gymnosperm genomes and *A. trichopoda* gene sets for single copy (light blue), fragmented (green), duplicated (light orange) and missing BUSCOs (burgundy) from embryophtya_odb10. **c,** Comparison of duplicated gene counts reported in gymnosperm genomes and *A. trichopoda*, whole genome duplications (light green), tandem duplications (dark green), proximal duplications (light orange), transposed duplications (light blue), single copy genes (burgundy). **d,** Repeat composition of *W.nobilis*1.0 genome.

Transcriptome profiling was performed in triplicate using polyA+ RNA isolated from three organs: leaves, ovules and pollen. In addition, one library from roots was also used for annotation purposes. Leaves and roots were collected from the same individual sampled for genome sequencing, while the ovule and pollen samples were collected from another clonal propagule. Over 20 million paired-end reads were generated from each RNA-seq library of which >85% mapped uniquely to the *Wollemia* genome (Supplementary Table S1.3). To evaluate the completeness of the assembly, BUSCO was run in genome mode against *embryophyta_odb10* of 1614 genes ^41^. BUSCO estimated the completeness of the assembly to be 90% which consists of about 1453 complete BUSCOs of which 1370 were single copy, 83 were duplicated, 97 fragmented and about 64 BUSCOs were missing (Supplementary Table S1.4). Completeness of other gymnosperm assemblies (Supplementary Table S1.2) range between 49 and 86%, suggesting *Wollemia nobilis* 1.0 genome completeness is better than assemblies of existing gymnosperm genomes ^42–45^. Despite the contiguity of the assembly, the BUSCO completeness of the *Wollemia* genome appears lower compared to annotated genes, likely due to the presence of very long introns in conifers, which can inhibit identification of genes ^45^ by BUSCO and other annotation tools. Indeed, the BUSCO completeness results are slightly higher within the annotated genes (see below). Further assessment of the *Wollemia* genome assembly was done using the alignment of Illumina DNA sequencing data, and the assembled transcripts from RNASeq data. To evaluate the coverage of the obtained genome assembly, high-coverage Illumina sequencing data and transcriptome reads were mapped against the assembled genome. The average mapping rate of RNA reads was 93.45%. (Supplementary Table S1.3)

### Annotation of *Wollemia nobilis* reference genome 1.0

Using a combination of RNA sequencing (RNA-Seq) based predictions, orthologs from other published gymnosperm genomes (**Methods**) and other open source transcriptomic data from closely related species from treegenesdb.org ^46^, we annotated 37,485 protein coding genes in the *Wollemia nobilis* 1.0 genome assembly. The mean coding sequence length and intron length of the annotated gene models were 1184 and 7745 bp, respectively (Supplementary Table S1.1). We annotated nearly five genes with extremely long introns above 1Mbp with the maximum intron length of 1.49 Mbp, one of the longest plant introns to date (Supplementary Table S1.2). Large introns are characteristic of conifer genomes, with introns up to 800 Kbp observed in *P. taeda* ^47^ and introns over 500 Kbp in *P. lambertiana ^48^*.

Of the 37,485 *Wollemia* gene models, 27,223 gene models were functionally annotated by either sequence similarity or gene family assignment with ENTAP ^49^, InterProScan5 ^50^ and TRAPID ^51^. Functional annotation revealed about 72.62 % of the genes had conserved domains and associated gene ontology terms (GO) (Supplementary Table S1.5). Gene sets were assessed with BUSCO in the protein mode against the Embryophyta orthodb10 set, which estimated the annotation to be 93% (Supplementary Table S1.4) complete with nearly 1500 complete BUSCOs, out of which 1415 BUSCOs were single copy (87.7), 85 were duplicated (5.3%) and 35 (2.2%) were fragmented models and the remaining 79 (4.58%) genes were uncovered (Supplementary Table S1.4). This indicates a higher quality of this genome annotation than for other published gymnosperm genomes, such as those of *Ginkgo biloba* (92.5%) and *Gnetum montanum* (88.1%) (Figure 2a). Further assessment of the gene annotation was performed using alignment of RNASeq reads with the genic regions of the assembly using qualimap. The average percentage of genes aligned to the exons, introns and intergenic regions was 74.9%, 12.5% and 13.32%, respectively (Supplementary Table S1.3). Across all RNAseq libraries, detectable expression (>1 CPM in ≥3 libraries) was observed for 18,132 genes (Supplementary Table S2.1). Of these, approximately 6,128 genes had highly tissue-specific expression (**τ** > 0.9 ^52^).

The 37,485 protein coding genes in the *Wollemia* genome were further analyzed to identify critical transcriptional, post-transcriptional and post-translational regulators with the goal of estimating the regulatory complexity in this ancient lineage. Specifically, the *Wollemia* gene models were processed to identify Transcription Factors (TFs), and small RNA processing components. Separately, small RNA populations were profiled from multiple tissues to estimate the full miRNA and siRNA complement produced in *Wollemia*. Clonal propagules of Wollemi pine only rarely produce male and female cones outside their climatic range, but samples of pollen and ovules were obtained from a single ramet at the Botanic Gardens in Padua, Italy. Sections of fixed material at the same stage revealed that the *Wollemia* ovules were harvested before fertilization, while pollen was harvested at maturity, comprising three haploid vegetative cell types, and two sperm cells (11 nuclei in total) (Supplementary Figure 3), as previously described ^53^. We also compared these *Wollemia* protein coding genes to seven other published genomes spanning the major land plant groups (**Methods**) and identified 13,923 orthologs shared with other species (Supplementary Figure 4).

#### Transcription factors and gene regulatory networks

To better understand the genetic basis of Wollemi pine architecture and reproduction, we investigated transcription factor (TF) families and small RNAs with a focus on those that may underlie *Wollemia*’s morphology and evolution. The complete gene models from the *Wollemia* genome (n=37,485) were scanned to identify all gene models that encode a transcription factor (TF) ^54^. The *Wollemia* genome encodes a total of 1,327 TFs including members from all 57 known plant TF families ^54^, with the exception of the FAR1 family. The FAR1 family is only present in the angiosperm lineage ^55^. Transcriptome profiling in this study is limited to the leaf, pollen and ovule tissues, but nonetheless detected robust expression (> 0.5 counts per million) of 710 TFs (Figure 3a).

**FIGURE 3.**
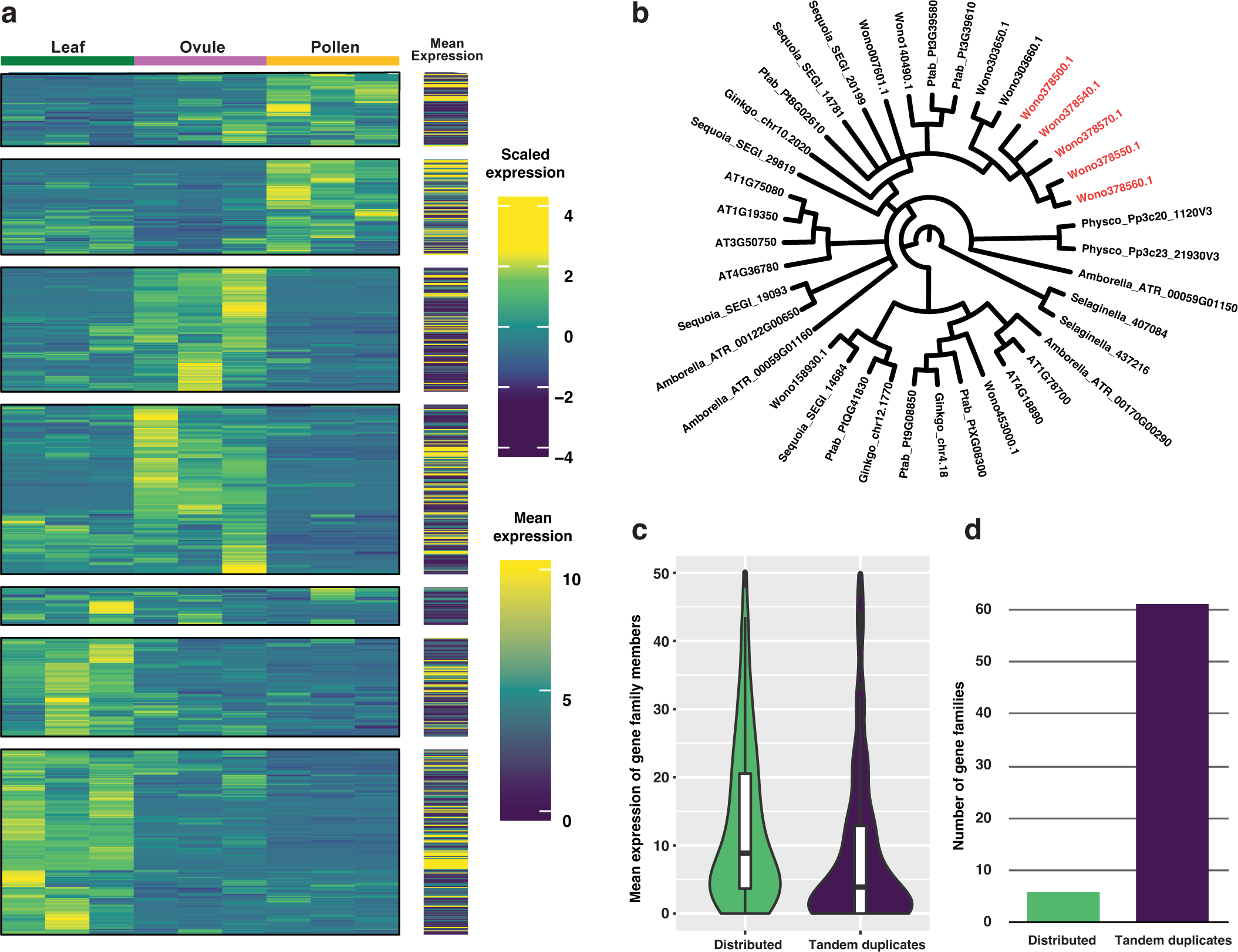
Tandem duplication expands gene families but majority of duplicated genes lack evidence of expression. **a,** Expression heatmap of all Transcription Factors (TFs) with detectable expression in at least one of the three sampled organs reveals organ-specific expression patterns for the majority of TFs. **b,** The family of BES1 (BRASSINOSTEROID SIGNALLING1) TFs is shown as an exemplar of TF families with relative expansion in the *Wollemia* lineage. Wollemi genome encodes additional BES1 orthologs primarily due to a cluster of tandemly duplicated copies in a single locus (in Red). However, none of these tandemly duplicated copies show evidence of expression in the three sampled organs. **c,** Gene families that included both distributed and tandemly duplicated members were assayed to estimate mean expression level of distributed vs. tandemly duplicated family members. The mean expression of tandemly duplicated members was lower than distributed members. Further, as shown in **d,** a much higher number of gene families included tandem duplicates with no observable gene expression.

Several TF families are found in some, but not all, gymnosperms consistent with the placement of *Wollemia*. WOX1, involved in lateral organ development, such as leaf and vein formation in angiosperms, has an ortholog in *Wollemia*. A similar ortholog is present in *Taxus chinensis*, though its function in gymnosperms is currently undetermined. KANADI 2 with other proteins determines abaxial leaf identity in *Arabidopsis* and has an ortholog in *Wollemia* and *T. chinensis*. The NZZ/SPL TF family, which includes SPOROCYTELESS, currently has no reported members in the published gymnosperm genomes, and has 0-2 members in most sequenced angiosperm genomes ^54^. Using the SPL and TIE transcriptional repressors from *Arabidopsis thaliana* as queries for homology searches in our high contiguity assembly of *Wollemia* allowed the identification of four genes (Supplementary Figure 5) that contain the SPL and EAR motifs and are therefore putative orthologs of the NOZZLE/SPOROCYTELESS TF that plays a critical role in reproductive development ^56^. We found similar homologs in other gymnosperm genomes using NCBI BLAST (**Methods**).

The relative proportion of TFs in each family were compared between *Wollemia* and the reference plant genome *Arabidopsis thaliana,* in order to identify TF families that have a >2-fold lower or higher proportion between the two species (Supplementary Table S6). The M-type MADS box and B3 TF families have proportionally fewer members in *Wollemia*, while the BES1, Trihelix, ERF, LBD and Leafy TF families have proportionally higher members in the *Wollemia* genome. The two TF families with the highest increase in membership are the LBD family, known to play a role in lateral root formation ^57^ and shoot branching ^58^, and the Leafy family, which has a single member in *Arabidopsis* that acts as a pioneer TF and is responsible for transition of the vegetative meristem into a floral meristem ^59^.

The BES1 TF family shows a specific expansion in the orthogroup that includes BEH1/2/3/4 TFs from *Arabidopsis* (Figure 3b), which is fueled by tandem duplications at one locus in the *Wollemia* genome (Figure 3b). However, transcriptome profiling detected no expression from the tandem duplicated copies of this BEH gene family. Extended analysis across all TF gene families that include tandemly duplicated genes (n=237), revealed that in 61 such families there was no detectable expression from tandemly duplicated gene family members. Conversely, gene family members that are distributed across the genome lack expression in only six TF families, implying a strong bias towards lack of expression of the tandemly duplicated gene copies. One explanation is that these tandem arrays are silenced pseudogenes.

The gene expression data was further used to initiate an ensemble network inference approach^60^ (see **Methods**) to generate gene interaction networks from the gene expression values. The full expression matrix of 24,583 *Wollemia* genes included 1,009 TFs, and the network inference approach was used to infer interaction edges between these TFs and all genes. The aggregated network included 24,581 nodes and was filtered to remove noisy edges via a backboning approach ^61^, to retain a sparse network of 645,758 network edges. This aggregated backboned network was further filtered to only retain the top 10% edges by rank (Supplementary Table S2). This high-stringency network now includes the higher confidence predictions of TF-target relationships in *Wollemia* and can be investigated to identify regulators of traits such as gamete development, pathogen response etc.

#### microRNAs target transcription factor and disease resistance genes

We performed small RNA sequencing from leaves, ovules and pollen, and used SHORTSTACK to separate small RNA into miRNA and siRNA populations, including transposon siRNA, tasiRNA (trans-acting small interfering RNA) and phasiRNA (phased secondary small interfering RNA) (Figure 4a). The small RNA length distribution in *W. nobilis* leaf, ovule, and mature pollen grains revealed a preferential accumulation of 21-nt sRNA in the leaf, while ovule and pollen additionally accumulated 24-nt sRNA (Figure 4b). Intriguingly, almost 50% of the leaf sRNA content is composed of 21-nt phasiRNAs (Figure 4a) which arise from 281 clusters (Extended Data Figure 1).

**FIGURE 4.**
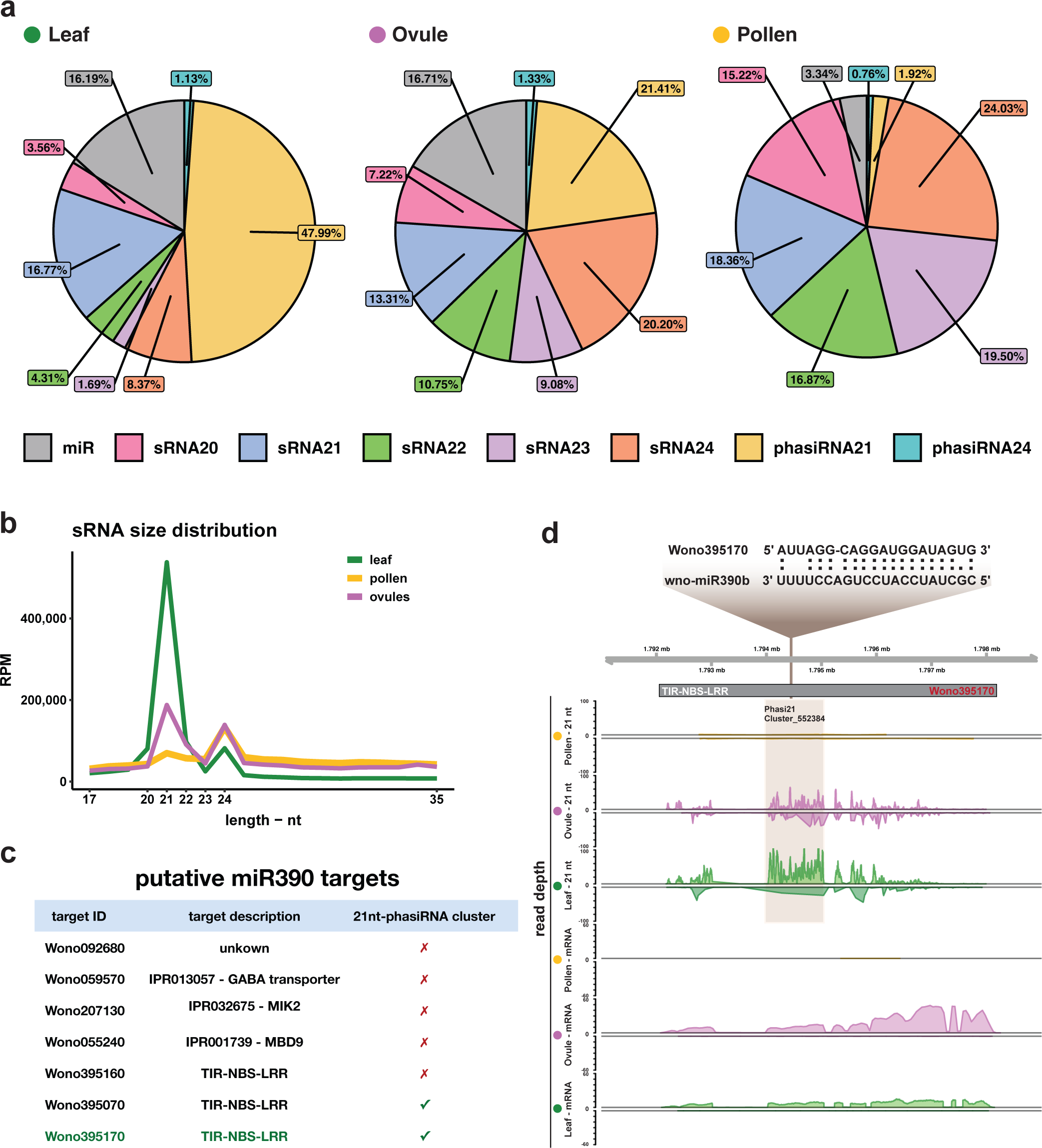
Non-canonical R gene 21-nt phasiRNAs accumulate in leavesand are triggered by miR390b. **a,** Small RNA classes in leaf, ovule, and pollen. **b,** Small RNA size distribution in leaf (green), ovule (purple), and pollen (yellow), revealing that leaf tissue accumulates more 21-nt small RNAs compared to other tissues, while ovule and pollen accumulate 24-nt small RNAs at the same level. **c,** wno-miR390 target prediction and overlap with 21-phasiRNA clusters. **d,** wno-miR390b, aligned with its putative target Wono395170 at the predicted target site. Read depth of the 21-nt phased siRNA (Cluster 552384), and Wono395170 transcripts.

Through a conservative approach (**Methods**), 277 microRNAs were identified, of which 38 were conserved (Supplementary Table S3.1). In common with angiosperms, Wollemi pine pollen grains accumulate more siRNA than miRNA, which comprise only 3.34% of total pollen sRNAs (Figure 4a, Extended Data Figure 1). We looked for transcription factors (TFs) among putative targets and found substantial conservation between canonical miRNA and their respective TF targets (Supplementary Table S3.2). For instance, both wno-miR156 and wno-miR535 families target *SPL-like* (*SQUAMOSA-promoter binding*) genes (Supplementary Table S3.2) ^62^ while the wno-miR166 family targets *HD-ZIP III* genes such as *PHB (PHABULOSA).* ^63,64^. In Arabidopsis, miR166 was reported to regulate the germ cell determinant *SPOROCYTELESS/NOZZLE* (*SPL/NZZ*), by targeting *PHB*, which in turn binds to the *SPL/NZZ* promoter to inhibit its activity ^65^. Our data suggest that in Wollemi pine, wno-miR159 also targets *SPL/NZZ* directly (Supplementary Table S3.2), as it does in tomato ^66^. As SPL/NZZ has so far not been found in other gymnosperms, these data suggest a novel pathway, conserved in some angiosperms and in *Wollemia*.

The Wollemi pine genome encodes the two canonical families of plant defense genes, namely the Defensins and the NBS-LRR genes. A total of 41 putative Defensin genes were identified by the presence of the conserved core Cysteine domain ^67^ – much less than the 300 genes computationally predicted in *Arabidopsis*. Defensin genes are arranged in multiple tandem clusters arising from gene duplication across the genome (Supplementary Figure 6e). NLR annotation predicted 697 putative loci, of which only 329 loci were confirmed to encode a complete NBS-LRR protein. Overlapping these 329 loci with the filtered gene model set yielded a set of 245 *Wollemia* gene models that encode a complete NBS-LRR gene. All subfamilies of NBS-LRR genes are represented in the genome with the largest subfamily being the TIR-LRR subfamily with 154 members ^68^ (Supplementary Figure 6d).

In many angiosperms and gymnosperms, NBS-LRR genes are targeted by the miR482/miR2118 superfamily of miRNAs giving rise to phasiRNAs in reproductive tissues, that depend on *RDR6* and *DCL4* ^69^. In some angiosperms and gymnosperms, additional miRNA target R genes in the leaves. In Wollemi pine, 260 NLR genes - 205 TIR-NLR, 6 CCR-NLR, 42 CC-NLR, and 7 CCG10-NLR - are potentially targeted by miRNA [SuppTable S3.3]. 31 miR482-like miRNA, ranging from 20 to 22 nt long, (Supplementary Table S3.4), target 51 of these NLR genes (Supplementary Table S3.4), of which 39 (30 are TIR) enclose phasiRNA clusters within their *loci* (SuppTable S3.5], and account for 15% of all expressed 21nt-phasiRNAs.

The TAS-miR390-ARF is another conserved pathway that produces 21-nt tasiRNAs that target ARF transcription factors and require *AGO7* as well as *RDR6* and *DCL4* ^70^. In the gymnosperm *Picea abies*, this pathway is conserved and includes several additional gymnosperm-specific miRNAs ^71^ Intriguingly, our data indicate an additional pathway in the Wollemi pine, where wno-miR390b also targets putative R-genes such as *MIK2* and three TIR-NBS-LRR genes (Figure 4c). In particular, one TIR-NBS-LRR gene (Wono395170) gives rise to abundant 21-nt phasiRNAs in both ovule and leaf (Figure 4c,d), where *DCL4* and *RDR6* are highly expressed (Extended Data Figure 1c,e). Unlike in *P. abies*, wno-miR390 accumulation is detectable in mature Wollemi pine leaves, where it accounts for 0.4% of R gene 21nt-phasiRNAs. Abundance in the ovule is higher, where it generates one tasi-ARF that potentially targets 8 ARFs (Supplementary Table S3.6). The *Wollemia* genome encodes multiple *AGO*, *DCL*, *CMT*, and *RDR* genes, representing each of the major clades found in angiosperms, which are differentially expressed in leaves, ovules and pollen, consistent with our small RNA analysis (Supplementary Figure 6g-j, Extended Data Figure 1).

Both R genes and microRNAs are dynamic components of the genome, and NLR genes are targeted by several different miRNA in gymnosperms, including *Picea abies* ^71^. Some miRNAs evolve from NLR duplicated genes in an “*evolutionary feedback loop*” ^72^. MicroRNAs modulate R gene expression, as the constant expression of resistance genes is detrimental to the host ^72^. The generation of secondary siRNA amplifies the regulatory signal but could also trigger silencing at secondary targets ^73^. During pathogen infection, *MIR* transcripts encoding miRNAs that target R genes are often down-regulated, accompanied by a decrease of phasiRNAs and an increase of R gene expression^73^.

### Transposable elements and gene silencing

The transposon annotation algorithm EDTA ^74^ predicted 63,908 unique transposable element-related sequences in the Wollemi pine genome, with widely varying copy numbers representing 86.18% of the genome (Figure 2d). Prominently, LTR retrotransposons (LTR-RTs) comprise 58.3% of the repeated TEs, and are further broken down into 18.94% *Ty3/Gypsy*, 14.22% *Ty1/Copia* and 25.14% unknown LTR. The unknown LTR-RTs are specific to *W. nobilis* and other gymnosperms. We computed the insertion times of intact LTR-RTs, by calculating the similarity between LTR, which are copied from each other during transposition. We detected a continuous accumulation of insertions from ∼33 MYA until now, with a burst around ∼5-6 MYA, as recently found in the genome of the Chinese Pine (*P. tabuliformis*) ^39^ (Extended Data Figure 2a). This reflects the high level of retention of ancient LTR retrotransposons in gymnosperms (low solo LTR ratio), in comparison with median TE insertion time ∼2.4 My BP in angiosperm genomes ^75^. non-LTR retrotransposons, and DNA transposons were also detected (Figure 2d). Interestingly, we found that the most frequent transposable element class in Wollemi pine are Helitrons, and that it had an unusually high number of DNA transposons overall (Figure 2d).

We assessed the number of active TEs in each class using transcriptome analysis in both somatic and reproductive tissues (ovules, pollen and leaves). Uniquely mapping reads were used to provide a conservative estimate of expressed TEs, which were then expanded to encompass nested elements and multi-mapping reads. We found thousands of transposons were highly expressed, especially in ovules. And while less than 2% of the members of each superfamily are active, the absolute number of active transposons (tens of thousands) likely reflects and is consistent with substantial transposition activity over time (Extended Data Figure 2 and Supplementary Table S4). The retrotransposon *Penelope*, showed the highest percentage of active elements in all three tissues analyzed 2.17%, 2.13%, and 4.07% in leaf, ovule, and pollen, respectively. *Penelope* is found in pines and other conifers, but not in other gymnosperms or angiosperms, and is thought to have transferred to conifers by horizontal transfer from an arthropod about 340 MYA ^76^. In absolute numbers of active elements (including nested elements and multi-mappers), both *Gypsy* LTR and *Mutator* TIR have close to 12,500 active elements in ovule, along with, 105 *Penelope*, 407 LINE and 5 other non-LTR (Supplementary Table S4).

DNA methylation plays an important role in regulating transposable elements, and can occur in three contexts in plants: CG, CHG, and CHH (H is A, T, or C) DNA methylation is reprogrammed to some extent in flowering plants during pollen development ^77^. This reprogramming is thought to help maintain genome stability and reset imprinted genes during reproduction ^78^. To investigate methylation levels in reproductive tissues, we performed whole-genome bisulfite sequencing (WGBS) in mature pollen grains and ovules. We found that the Wollemi pine genome is highly methylated at very similar levels in pollen and ovules in all contexts: CG 77.5% and 78.4%, CHG 68.8% and 65.05%, CHH 2.7%, and 2.5% respectively (Extended Data Figure 3). Transposable elements are highly methylated in both CG and CHG contexts but they are less methylated in mature pollen grains than ovules (Extended Data Figure 3). Importantly, actively transcribed elements are robustly demethylated in both tissues relative to silent elements (Figure 5a-f). Most notably, active *Penelope* elements completely lose CHG methylation in pollen and in ovules, where they are preferentially expressed (Figure 5e-f). CHG methylation is closely correlated with histone H3 lysine 9 methylation in plants, suggesting that *Penelope* elements may be regulated by this epigenetic modification in the Wollemi pine germline. Remarkably, *Penelope* elements in Drosophila undergo similar regulation in ovaries ^79^.

**FIGURE 5.**
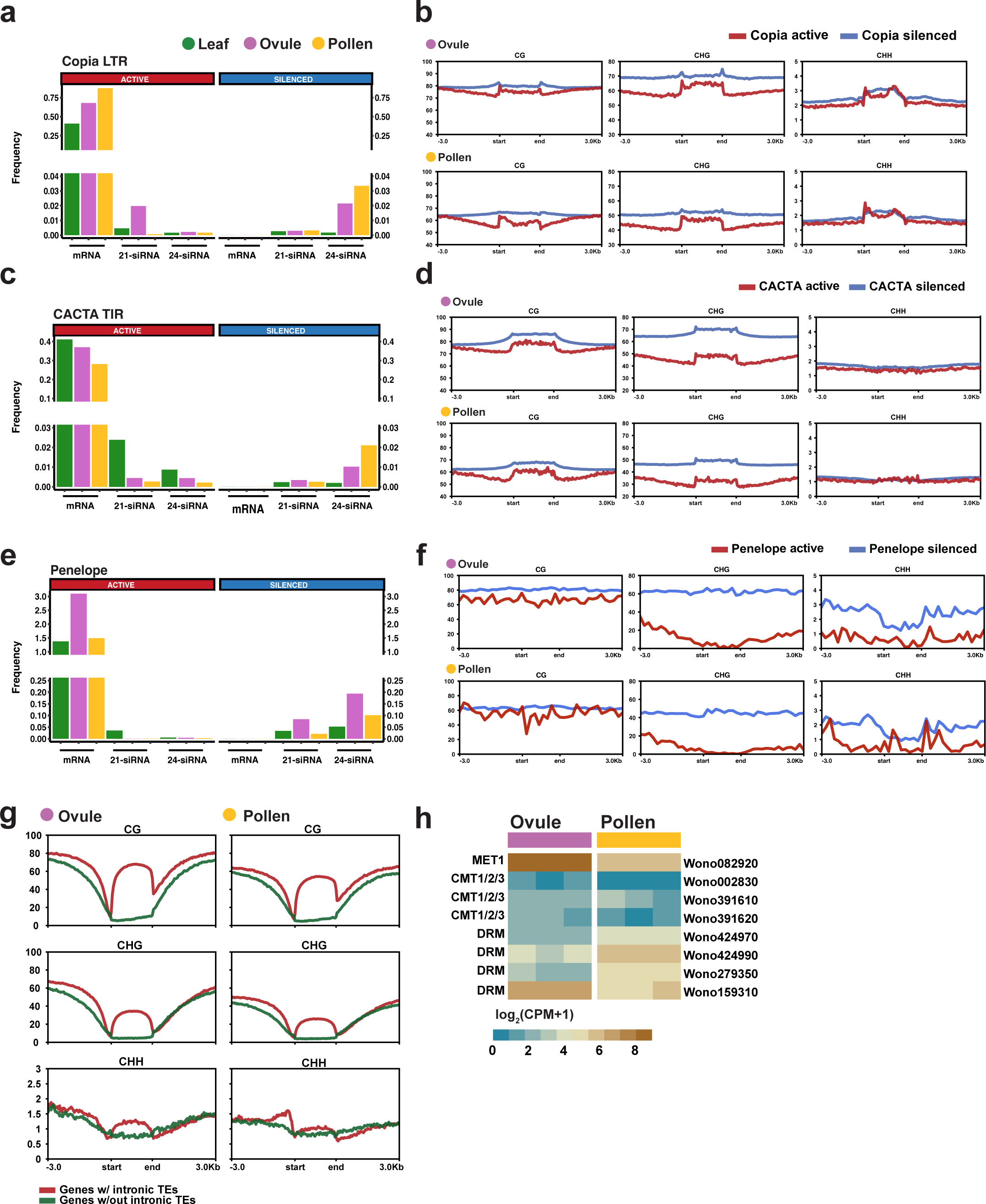
Active *COPIA*, *CACTA* and *Penelope* retrotransposons are silenced by DNA methylation and siRNA. **a,** mRNA and small RNA sequencing of *Copia* LTR retroelements.CPM per transposon copy (frequency) is shown for leaf, pollen and ovule sequencing reads. Silent elements generate 21 and 24-nt siRNAs in ovules and pollen grains, while expressed elements generate mRNA transcripts instead. **b,** Metaplots of methylation patterns in active and silenced *Copia* LTR retrotransposons in each sequence context, CG, CHG and CHH. CG and especially CHG methylation decreases in actively transcribed TEs. **c-d,** Similar patterns of small RNA and transcript accumulation as well as DNA methylation were found for active and silent *CACTA* TIR DNA transposable elements in ovule and pollen grains. **e,-f,** Active *Penelope-like* elements had the most abundant transcripts in ovules and pollen grains, while silent elements generate 21 and 24-nt siRNAs. Both CG and CHG methylation were lost from active TEs compared with silent TEs. **g,** DNA methylation in gene-bodies with (red) and without (green) intronic TEs. **h,** *MET1* (*Methyltransferase1*), *CMT1/2/3* (*Chromomethylase1, 2, 3*), and *DRM* (*DOMAINS REARRANGED METHYLASE*) expression levels were compared in ovules and pollen.

We also found that the genes containing intronic transposable elements (20,388 genes out of 37,484 annotated genes) are methylated in both CG and CHG contexts, while the genes without elements within introns are not methylated in any context (Figure 5g). *P. tabuliformis* (Chinese pine) has a very similar methylation profile in both TEs and genes containing TEs in shoot tissue ^39^ as does *P. abies* (Norway Spruce) in leaf (LF) and during somatic embryogenesis (SE) from cultured cells ^80^. These observations are consistent with the idea that methylation in introns is restricted to TE, and that there is no gene body methylation in Wollemi pine and likely other conifers. *Welwitschia mirabilis* has similar CG and CHG methylation patterns, but CHH methylation is significantly higher, 35.7%, in both meristem and leaf ^39^. The reduction of CG and CHG transposon methylation in Wollemi pine pollen suggests reprogramming may occur as it does in angiosperms, despite the lack of double fertilization ^77^.

In most angiosperms, CHH methylation depends in whole or in part on RNA directed DNA methylation (RdDM). The RdDM pathway has been a controversial topic in gymnosperms, because of the lack of 24nt siRNA in leaf tissues which has been widely observed. However, in gymnosperms and other non-flowering plants, most genes required for a functional RdDM pathway have now been found ^80–82^. In *Wollemia*, we found (Extended Data Figure 5) four *DCL3* homologs, required for 24nt siRNA, as well as all the other RdDM genes essential for a functional pathway, indicating canonical and non-canonical RdDM pathways are active (Extended Data Figure 5). Leaf tissue had the lowest levels of 24-nt TE-siRNA, accompanied by very low expression of *RDR2* which is also required for their production [Extended Data Figure 5]. By contrast, abundant 24-nt siRNA were found in ovules and pollen, accompanied by expression of *RDR2*. 24nt siRNAs corresponded to silent TEs, while actively transcribed elements did not accumulate them (Extended Data Figure 2), as illustrated by the *Copia* LTR and *CACTA* TIR superfamilies (Figure 5a,b,c,d and Extended Data Figure 4). The silenced elements also had higher methylation levels, especially in the promoter and terminal repeats and in the CHG context. Expression levels of *MET1* (*METHYLTRANSFERASE1*) and one out of the four *DRM* (*DOMAINS REARRANGED METHYLASE*) homologs were higher in ovule when compared to pollen, while the other three *DRM* homologs were increased in pollen grains. (Figure 5h).

#### Population structure and genetics

We re-sequenced the genomes of 5 Wollemi pine individuals, collected in Wollemi National Park from each of four closely located canyons. Read coverage was 21× for sample NSW1032268, 15x for NSW1032275, 14× for NSW1032290, 16× for sample NSW1032313, and 72× for sample NSW1023645. We found a total of 12,576,749 variants for all genomes. The number of SNPs for each *Wollemia* genome ranged from 2,074,876 to 6,864,461. Nucleotide diversity was low, ranging from 4.53E-04 to 5.63E-04 (Supplementary Table S5.1). The absence of homozygous variants was consistent with a preponderance of somatic variants in clonal populations, although there was a high frequency of missing data in these very large repetitive genomes, ranging from 0.45 to 0.83 (Supplementary Table S5.1).

We also performed targeted genotyping of unmethylated regions of the genome using DArTseq^83^, from 99 trees and saplings derived from approximately 60 individual ortets comprising the extant population. The filtered DArTSeq dataset contained 14,118 SNPs across 99 samples, reduced from 20,809 raw SNPs. Low genetic variation was found in the DArTSeq SNPs, where 66% of the filtered sites were monomorphic and 4,710 sites were variable. The final dataset had a 25.7% rate of missing data. The clustering analysis revealed four genetic clusters (Figure 6a-c), which corresponded to the sampling locations of the individuals. Low admixture among all genetic clusters was observed (Figure 6a) with fixed genetic differences between all but one population pair (S1-S4, which also have the closest geographic locations), ranging from 29 to 94 fixed SNPs per pairwise population comparison (Supplemetary Table 5.2), and high pairwise F_ST_ values, ranging from 0.44 to 0.58 (Supplementary Table S5.2). Observed heterozygosity (5.3-8.1×10^-6^) was lower than expected heterozygosity (6-15×10^-6^) (Supplementary Table S5.2).

**FIGURE 6.**
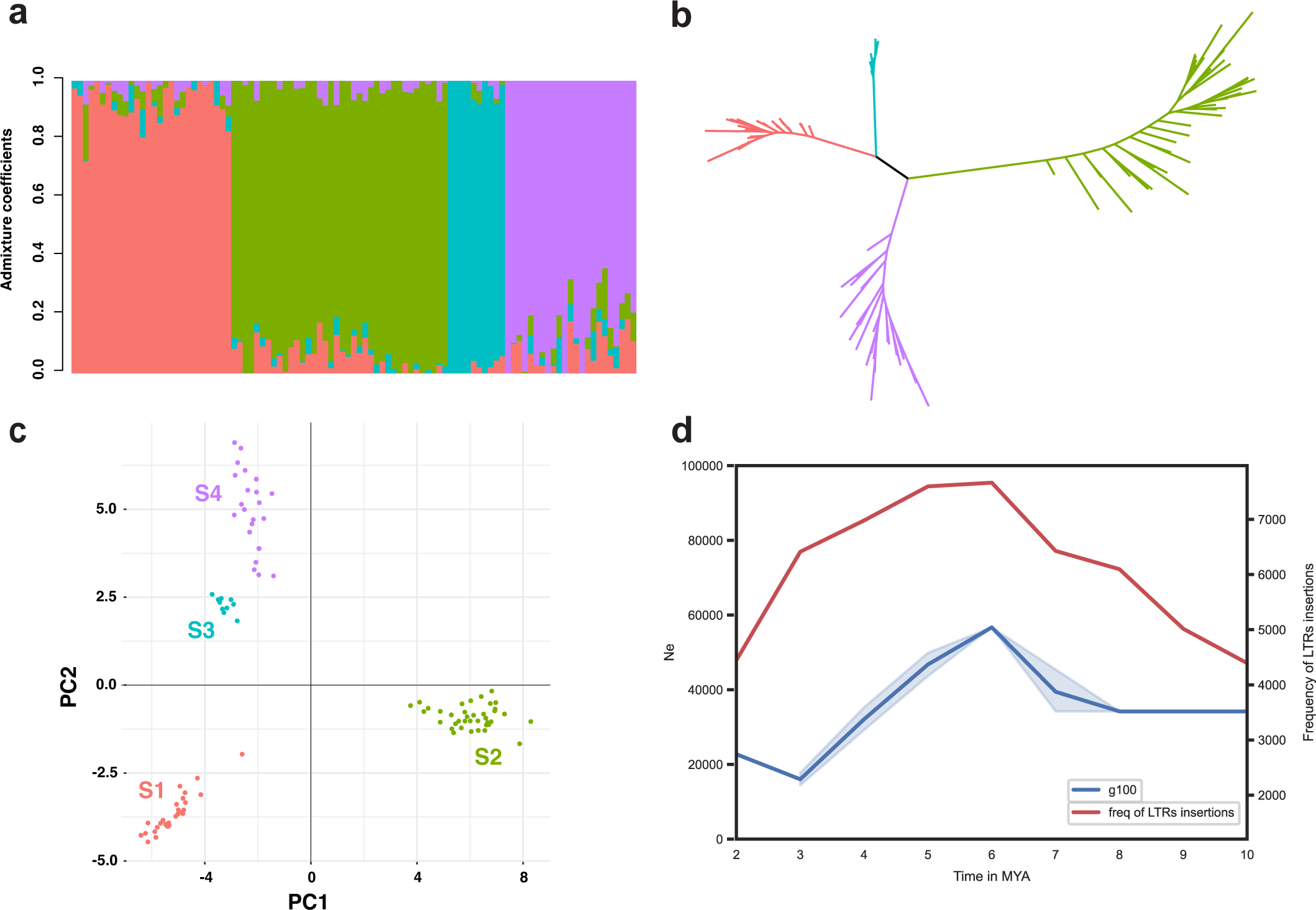
Population genetic structure and historical demography of *W. nobilis*. **a,** Population cluster assignment based on DArTSeq analysis and admixture coefficients (y axis) for 99 samples from approximately 60 individuals from four nearby locales (bars). The colors correspond to the population assignments of the cluster analysis (S1, red; S2, green; S3, blue; S4, purple). **b,** Unrooted phylogeny of *W. nobilis* samples based on DArTSeq *loci*. **c,** Principal Component Analysis plot showing genetic similarity and distances between samples. **d,** Population size change for 4 full genome-sequenced individuals representing each genetic cluster (blue) and LTR retrotransposon insertion rate variation through time (red).

We reconstructed changes in *Wollemia* population size through time by implementing a sequentially Markov coalescent model based on the site frequency spectrum of the *Wollemia* whole genomes ^84^, and assuming generation times of 100, 30 or 10 years. We found very similar *Wollemia* population size through time curves for all individuals, although they were sampled in different locations corresponding to the four populations identified with the SNP dataset. The *Wollemia* populations exhibited a steep increase in size from 6-8 My BP, 1.5-2.5 My BP or 550-800 ky BP, assuming generation times of 100, 30 or 10 years, respectively (Figure 6d, Supplementary Figure 7). The increase was followed by a continuous period of decrease to around one-fifth of the ancestral size, spanning most of the recovered demographic history, from 7-3 My BP, 2.2-0.8 My BP and 750-300 ky BP for generation times of 100, 30 and 10 years, respectively (Figure 6d, Supplementary Figure 7). The final event recovered was a slight population size increase around 3 My BP, 800 ky BP and 300 ky BP for generation times of 100, 30 and 10 years, respectively. There was a marked overlap between the peak and decrease of insertion rate of transposable elements (LTRs) and the demographic curves based on a generation time of 100 years. This generation time is in line with araucarias^85^. It is likely that the *Wollemia* populations underwent steady decline in size during the transition from the late Miocene to the Quaternary period.

In addition to the whole genome based demographic reconstructions, we tested different demographic models for each of the four *Wollemia* populations by implementing a composite-likelihood approach based on the DArTSeq dataset. Population demographic modeling yielded bottlenecks as the most likely scenario for all but one population, for which the decrease model had the best fit (population S1, Supplementary Table S5.3). However, the point estimates of the parameters for the bottleneck models were not contained within the confidence interval in many cases (e.g. population S2, Supplementary Table S5.4), suggesting an overall poor fit of the bottleneck model to the data. The estimates of time of population size decrease or onset of population bottleneck overlapped for the four populations, ranging from 10-26 ky BP (Supplementary Table S5.4). The best joint demographic history model consisted of an ancestral population size decrease followed by split into daughter populations S1 and S2 (Supplementary Figs. 7 and 8), with a posterior probability of 82.3% (Supplementary Table S5.5). The point estimate for ancestral decrease in population size was 53,600 generations BP (95%CI: 10,700-97,100; Supp Table 5.6). However, the posterior distributions of the time of population size decrease largely overlapped prior ranges (10^3^-10^5^ generations with a uniform prior distribution, Supplementary Figure 7), and the posterior normalized error was high for this parameter (0.9, Supplementary Table S5.6). A larger SNP dataset is likely needed to recover ancient changes in population size such as the late Miocene/early Pliocene decrease recovered with the whole-genome approach. The point estimate for the split between populations was 14,700 generations BP (4.5-25.1K 90% credibility interval, Supplementary Figure 7).

## Discussion

In the late Eocene, the detachment of the Sahul continental plate from the southern supercontinent Gondwana caused the establishment of circumpolar currents and the drying and cooling of global climates. As Sahul moved further north, it progressively lost much of the broad-leaved vegetation types that once dominated ^86^. Using an average generation time of 100 years, our analysis suggests that the ancestral *Wollemia* population increased rapidly in population size around 8-6 my BP during the general drying trend as Sahul moved north. During this time, rainforest retreat and the expansion of open forest may have created a niche for an emergent such as *Wollemia* to expand its range. During the Pliocene (5.3-2.5 My BP) rainforests contracted further, becoming mostly confined to the east coast of Australia ^87^. Our *Wollemia* population data presented herein highlights a population shrinkage to one-fifth of the original size at 7-3 My BP, overlapping this aridification event. Presumably, further drying shrank forested areas of inland Australia and *Wollemia* was forced into a narrowing band of intermediate rainfall, near the Great Dividing Range. With the onset of the climatic fluctuations of the Quaternary (in the last 2.5 My BP), repeated cycles of contraction in response to the cool-dry glacial periods, and expansion in response to the warm-wet interglacials changed the distribution of many Gondwanan lineages ^88^. The Last Glacial Maximum was particularly intense, with high aridity and increased fire intensity ^89,90^ leading to extreme rainforest contractions and the replacement of araucarian-dominated forests by pyrophytic sclerophyllous vegetation, and a substantial loss of Gondwanan rain forest lineages ^91,92^. Studies exploring rainforest assembly patterns and multispecies landscape genetics have identified high levels of dynamism ranging from continental biotic exchanges to species level responses to changing conditions ^4,88,93,94^.

Species that were less capable of responding to cyclical changes in habitat availability and were constrained by biotic and abiotic factors, rapidly lost much of their original distribution and eventually became extinct or became significantly bottlenecked ^95–97^. *W. nobilis* is one such species with an extremely restricted range, associated with narrow, steep-sided sandstone canyons and gorges protected from fire and dominated by warm temperate rainforest ^98^. Less than 60 individuals reduced into four spatially proximal stands remain that are impacted by a range of soil borne pathogens including *Phytophtora cinnamomi* ^16^.

Changed and changing environmental conditions are also likely to impact the viability and limit distributional potential ^99,100^. Population sample sequencing reported here demonstrates expected high levels of inbreeding and clonal propagation with an extremely low heterozygosity, even for a gymnosperm genome. There was scant evidence of genetic transfer between the four main population groups separated by gorges that we sequenced. Wind direction and deep gorges limits pollen transfer and seeds are large and often eaten by cockatoos.

The genomic analysis of *Wollemia* pine presented here provides some clues as to the developmental and genomic features that governed population trajectories of a single species subject to extreme climatological changes over geological time. While our initial analysis is necessarily preliminary, availability of the sequence and related functional datasets will allow detailed exploration of these features in the future.

A preliminary study of the genome identified orthologs of angiosperm flowering genes, most also found in other gymnosperms. Most “flower” developmental genes are present in *Wollemia*, and many orthologs of these genes regulate cone development in other gymnosperms. Angiosperms clearly have expanded floral developmental genes, but developmental genes for sexual organs predate the gymnosperm/angiosperm split. Gymnosperms lack sepals and petals, and there is no *Wollemia* ortholog of APETALA1. But APETALA2, an A-function gene in *Arabidopsis*, is present as an ortholog in *Wollemia* (Supplementary Figure6) and is also found as two homologs in *Pinus thunbergii* where it is expressed in female cones ^101^. The B-function gene APETALA3 is not present as an ortholog but the B-function PISTILLATA has an ortholog in *Wollemia*, and is known in gymnosperms to regulate microsporophyll development ^102^, while in *Arabidopsis* PISTILLATA is involved in the analogous petal and stamen differentiation ^103^. APETALA3 and PISTILLATA are hypothesized to be derived from a duplication event early in the angiosperm lineage, and in gymnosperms are present as a single gene ^104^. An *AGAMOUS* ortholog is present in *Wollemia*, and as observed in other gymnosperms, this C-function gene presumably regulates development of reproductive organs. *NOZZLE*/*SPOROCYTELESS* functions downstream of *AGAMOUS* in reproductive organs and is also present, and presumably regulates sporogenesis and ovule development in *Wollemia* as in flowering plants ^105,106^. In Arabidopsis, this gene functions as a transcriptional repressor in ovule development, by repressing TCP class transcription factors, which are also involved in axial branching ^107^. Along with the observed expansion of LBD transcription factors, additional regulation of NZZ by miR159 in Wollemi pine may contribute to the presence of axillary buds in Wollemi pine and other Araucariaceae, unlike other conifers ^28^.

Developmentally important small RNA are also largely conserved in Wollemi pine, including many miRNA, tasiRNA and phasiRNA. While the genome sequence alone cannot reveal the function of individual genes, it is notable that fully half of the small RNA in leaf are phasiRNA targeting disease resistance genes (R genes). This is partly because of the unusual targeting of these genes by miR390, which normally targets the non-coding RNA *TAS3*, an important regulator of leaf polarity ^108^. It is conceivable that phasiRNA targeting of R genes in leaves and ovules could contribute to disease susceptibility in the Wollemi pine. The expression of miR390 in leaves is likely associated with the flat expanded leaves found in Wollemi pine, rather than the needles found in *P.abies* and most modern conifers, which have less pronounced polarity. It is possible therefore that regulation of leaf shape may have led inadvertently to reduced pathogen resistance.

Wollemi pine has abundant siRNA, although, in common with other gymnosperms, leaves are depleted of 24nt siRNA, unlike ovules and pollen. We have found this is due to the lack of expression of RDR2 in leaf tissues. This relaxed silencing regime in somatic tissues could result in transpositions that are subsequently inherited, as the germline arises constantly from the soma in long lived trees, resulting in progressively divergent germ cells over time ^109,110^. On the other hand, the correlation of 24nt siRNA with silent transposons demonstrates the conservation of transposon silencing pathways, including RNA directed DNA methylation, in the most ancient extant ovules and pollen analyzed to date. Intriguingly though, tens of thousands of transposons remain active in Wollemi pine, including hundreds of *Penelope* retrotransposons, which are thought to have transferred horizontally from arthropods to conifers more than 300 my bp ^76^. This is close to the estimated divergence time of the Araucariaceae from other conifers ^23^, and it is interesting to speculate that arthropod infestation may have played a role in their origin. Retention of *Penelope* activity in Wollemi pine suggests remarkable resilience to silencing mechanisms in this lineage.

Our population genomic analysis has revealed the highly inbred, and likely clonal relationship of the few remaining wild Wollemi pines consistent with their critically endangered status. This analysis has revealed reductions in population size that coincided with drier and warmer climates over geological time. This trend has rapidly accelerated in recent years in the face of man-made climate change, as sharply illustrated by the recent destructive bushfires that took a toll on the remaining stands. The expansion of repetitive, mostly transposon derived, sequences may have contributed to low population genomic diversity of *Wollemia*. A burst of retrotransposition seems to have coincided with population increase 6 my bp, followed by a sharp decline, but many transposons remain intact and expressed in the Wollemi pine genome, especially in reproductive tissues. We considered several scenarios consistent with this observation. First, expansion of transposon copy number may have contributed to subsequent population decline, associated with accumulation of deleterious insertions. Second, transposon expansion is favored by sexual reproduction ^111^, and if population decline was accompanied by a switch to clonal propagation, this may have reduced subsequent transposon activity. Finally, transposon activation, through unstable epigenetic suppression in mosaic reproductive tissues, may have increased epigenomic and genomic diversity during long generation times, and mitigated declining population size and associated lower fitness. Many organisms with low genetic diversity have very large genomes, *Wollemia* included, while organisms with high genetic diversity, such as angiosperms, often have smaller genomes with fewer transposons supporting this idea. The Wollemi pine has been rescued from extinction temporarily by propagating and distributing these highly inbred individuals around the world as clones, and further analysis of the genomes of their progeny may reveal clues to ensure their preservation.

## Methods

### Sample Collection [Reference Genome Sequencing]

*Wollemia nobilis* leaves were collected from individual 692/2013A*, grown in cultivation in the Nolen Greenhouse of the New York Botanical Garden (NYBG). The individual was acquired by the NYBG in 2013. Leaves were cut and immediately flash frozen in liquid nitrogen. Plant material was stored at −80 degrees C in the Plant Research Laboratory at the NYBG.

### Sample collection [Population Genomics]

Preserved leaf tissue from four individual *Wollemia* trees representing each gorge, and a further individual was sourced from the ex-situ *Wollemia* collection at the Australian Botanic Garden (ABG), Mount Annan west of Sydney (New South Wales, Australia). The leaf tissue was frozen at −80° for 48 hours, freeze-dried and preserved with silica gel preparatory to whole genome sequencing. All four sites fall within an approximate radius of 3.5km. Sites 1 and 4 with less than one kilometre between them are the closest in proximity.

Sampling for the DArT-seq population study aimed to represent as many individuals from the four sites as possible. Fresh leaf material for 87 samples was collected from several trips to the wild sites. Twelve samples (with known wild provenance) were sourced from the living collections at the Australian Botanic Garden, Mount Annan. These samples represent all accessible individuals from the wild including 23 distinct individuals from site 1; 33 from site 2; 5 from site 3 and 22 from site 4. An additional sixteen samples (5 from each of sites 1,2 and 3 and 1 from site 4) were sampled where it was unclear whether stems close together belonged to the same tree or different individuals or where there was a high amount of missing individuals. Leaf tissue was frozen at −80° before being freeze dried and stored in airtight containers with silica gel. Five to ten mg of leaf tissue from each sample were randomly assigned to plates and sent to Diversity Arrays Technology Pty Ltd for DNA extraction and DArTseq genotyping.

### Collection of ovules and pollen from cones of *Wollemia nobilis*

Plant material was collected during the first week of April 2021, from a fertile *Wollemia* plant cultivated at the Botanical Garden of Padova, Italy. Female cones were sampled and dissected to collect ovules under a dissecting microscope (Leica EZ4W). Ovules were collected in 1.5 mL plastic tubes and immediately frozen in liquid nitrogen. Male cones were placed in 50 mL plastic tubes and then shaken using a vortex to release pollen in the tube. After shaking, tubes were briefly centrifuged (1 minute at 3000 rpm) to collect pollen at the tube bottom. Cones residues were removed, and pollen was immediately frozen in liquid nitrogen. All the samples were stored at −80°C until use.

### RNA extraction

RNA extractions were performed in triplicate: for *Wollemia* ovules, each replicate consists of ovules from two cones pooled together; for pollen, each replicate consists of pollen collected from one pollen cone. RNA was extracted using the protocol described by Chang *et al.* ^112^. RNA was resuspended in 20 μL of RNase-free water. For each sample, an aliquot of approximately 10 μg of RNA was treated with 2 units of DNase (NEB) in a final volume of 100 μL for 10 min at 37°C. The reaction mixture was purified using Zymoresearch RNA purification kit and clean RNA was eluted in 17 μL. Sample purity and concentration were evaluated using a NanoPhotometer® (IMPLEN).

Leaf RNA extractions were performed in triplicate, pooling leafs from different branches of the same tree at the NYBG using PureLink™ Plant RNA Reagent (Invitrogen) following the manufacture’s recommendations, followed by DNase treatment, using Turbo DNase™ (Invitrogen). The RNA quality was evaluated with Bioanalyzer (Agilent), Agilent RNA 6000 Nano kit.

### DNA extraction

Pollen from two *Wollemia* cones was pooled and then divided in three parts and extracted separately; ovules from two cones were pooled and then subdivided in three parts and extracted separately. DNA extractions were performed following the protocol described by Doyle ^113^, with the addition of 2% of PVP in the extraction buffer. Extracted DNA was resuspended in 20 μL of DNase-free water. DNA concentration and sample purity were evaluated using a NanoPhotometer® (IMPLEN).

### High Molecular Weight DNA Extraction

#### Nuclei Isolation

High Molecular Weight DNA extraction from young leaves of a clonal propagule of Wollemi pine at the New York Botanical Garden, was performed according to the Circulomics Nanobind Plant Nuclei Big DNA Kit (NB-900-801-01). First, Nuclei were isolated using the protocol developed by Workman et al. ^114^. Three solutions were prepared for use in the Nuclei Isolation Buffer: 10X Homogenization buffer stock *(100 mM Trizma base, 800 mM KCl, 100 mM EDTA, 10 mM spermidine trihydrochloride, 10 mM spermine tetrahydrochloride)*; 1x HB solution *(1× HB, 0.5 M sucrose)*, 20% Triton X-100 stock *(1× HB, 0.5 M sucrose, 20% (vol/vol) Triton X-100)*. For the final Nuclei Isolation Buffer, to be prepared the day of the extraction: 1× HB solution, 0.5% Triton X-100 solution, 0.15% (vol/vol) 2-mercaptoethanol.

For each DNA extraction, 1 gram of young leaf tissue was ground in liquid nitrogen. Cells were lysed by placing plant material with 20 ml of Nuclei Isolation Buffer in a 250 ml bottle. All steps of nuclei isolation were performed at 4°C. The bottle was placed in a 2 L beaker filled with ice, and the lysate stirred for 20 minutes at high speed on a stir plate. The homogenate was then gravity filtered through 5 layers of Miracloth into a 50 mL tube and centrifuged at 1900 x g for 20 minutes. The supernatant was discarded and the nuclei pellet was resuspended gently using a small paintbrush and cold Nuclei Isolation Buffer. The nuclei were centrifuged again at 1900 x g for 10 minutes. This wash process was done 3 times, or until there was no visible color remaining in the nuclei pellet. After the final wash, the nuclei were resuspended in 1 mL HB buffer and transferred to a 1.5 ml Protein LoBind microcentrifuge tube. The nuclei suspension was pelleted at 5000 x g for 5 minutes and flash frozen in liquid nitrogen until the nuclear DNA isolation step.

### Nanobind-assisted DNA Purification

Continuing with the Circulomics Nanobind Plant Nuclei Big DNA Kit protocol: frozen nuclei from *Wollemia* leaves were resuspended in 60 microliters of Proteinase K solution, and vortexed thoroughly until fully resuspended. Nuclei were treated with RNAse A, then lysed using buffer PL1 and thorough vortexing. Nuclei were then incubated for 30 minutes on a thermomixer set at 55 °C and 900 rpm. Mid- lysis pulse vortexing (5x) was performed to enhance mixing. Once lysis was complete, the lysate was centrifuged at 16000 x g for 5 minutes at room temperature. From this point forward, the DNA was handled delicately to prevent shearing. The supernatant was transferred to a new 1.5 mL Protein LoBind microcentrifuge tube using a wide bore pipette. The Nanobind disk was added to the supernatant followed by 1X volume of isopropanol. The solution was mixed slowly, on a HulaMixer for 20 minutes at room temperature. *(HulaMixer settings: Rotation 9 rpm; Tilting: 70°, 12s; Vibration: 2°, 1s).* Tubes were placed on a magnetic rack, and precipitated DNA was carefully washed 3x with buffer PW1. DNA was eluted from Nanobind Disk in 200 microliters buffer EB.

### Oxford Nanopore Libraries

Extracted DNA from *Wollemia* leaf nuclei was run on Femto Pulse to assess fragment length distribution. DNA was sheared to ∼50-75kb using a Diagnode Megarupter or via Needle shearing following manufacturer’s recommendations. The Circulomics short read eliminator XL kit, which iteratively degrades short fragments, was employed to enrich long DNA fragments to increase assembly contiguity. DNA was prepared for Nanopore sequencing using the ONT 1D sequencing by ligation kit (SQK-LSK109). Briefly, 1-1.5ug of fragmented DNA was repaired with the NEB FFPE repair kit, followed by end repair and A-tailing with the NEB Ultra II end-prep kit. After an Ampure clean-up step, prepared fragments were ligated to ONT specific adapters via the NEB blunt/TA master mix kit. The library underwent a final clean-up and was loaded onto a PromethION PRO0002 flow cell per manufacturer’s instructions. The flowcells were sequenced with standard parameters for 3 days. Basecalling was performed with Guppy V4 to increase quality. The genome was sequenced to ∼60X coverage, with 30X coverage in reads >= 30kb.

### Illumina Libraries

Illumina short read DNA libraries were prepared from the *Wollemia* leaf tissue with the Illumina TruSeq DNA kit, targeting a 550bp insert size with PCR enrichment. Libraries were sequenced at Texas A&M,, on a NovaSeq S4 flowcell in paired end 150bp format to ∼30x genome coverage.

### Transcriptome sequencing

RNA Seq libraries from *Wollemia* tissues (leaves, roots, ovules & pollen) were performed using the Kapa mRNA Hyper prep kit with 100ng total RNA in 50uL DNase free water. The mRNA was captured with oligo-dT beads and then fragmented using heat and magnesium. Fragmentation was performed for 6 minutes at 85C (selecting fragments of 300-400bp). The 1^st^ cDNA strand was synthesized using random priming. Then this is followed by a combined 2^nd^ strand synthesis and A-tailing. Illumina adaptor ligation was performed followed by library amplification using high-fidelity, low bias PCR (PCR enrichment of 13 cycles). The strand marked with dUTP was not amplified, allowing strand specific sequencing. QC for the final libraries was done using Qubit and Bioanalyzer. Samples were then pooled and the final library was quantified using Kapa qPCR. The pool was sequenced on a paired end 150bp high output NextSeq500 run.

### Whole Genome Bisulfite Sequencing

Genomic DNA from mature pollen and ovule DNA was isolated using CTAB buffer (2% CTAB, 100 mM Tris-HCl pH 8, 20 mM EDTA, 1.4 M NaCl, 2% PVP). For the bisulfite conversion, EZ-DNA Methylation Gold kit (Zymo Research) was used according to the manufacturer’s instructions, the converted DNA was then used for the library preparation using the NEXTFLEX® Bisulfite-Seq kit (Perkin Elmer) according to the manufacturer’s instructions. Each library was sequenced on the NextSeq high output platform, paired-end 150-nt. Mapping and methylation calls were processed using Bismark v0.23.1 ^115^. Downstream analyzes were performed using *deepTools* ^116^.

### Small RNA sequencing, annotation and mapping

Total RNA from leaf (4 technical replicates), mature pollen (3 technical replicates) and, ovules (3 technical replicates) was isolated using CTAB buffer (2% CTAB, 2% PVP-K30, 100 mM Tris-HCl pH 8.0, 25 mM EDTA, 2 mM NaCl, 0.5 g/L spermidine, 2% ß-mercaptoethanol). NEXTFLEX® Small RNA-seq kit v3 (Perkin Elmer) was used according to the manufacturer’s instructions to prepare the small RNA libraries. The barcoded samples were pooled and sequenced on the NextSeq high output platform, single-end 50-nt. For the small RNA annotation, the reads were pre-processed by trimming the adaptors using Cutadapt 3.4, merging the sample files (leaf, ovule, and pollen), and filtering by length (18 to 26 nucleotides). The filtered file was used as input for miRNA and siRNA annotation using ShortStack 3.8.5 (--mismatches 0 --foldsize 1000 --mincov 2) ^117^. The mature miRNA fasta file generated by ShortStack was blasted against *Aquilegia caerulea*, *Arabidopsis thaliana*, *Amborella trichopoda*, *Brachypodium distachyon*, *Oryza sativa*, *Picea abies*, *Pinus densata*, *Pinus taeda*, *Selaginella moellendorffii*, *Zea mays* mature miRNA sequences retrieved from miRBase 22.1 using BLAST+/2.9.0. Only the miRNAs with up to 1 mismatch and minimum length identity of 19 nucleotides were considered We used fastsimcoal2 v2.7 ^130^ to test demographic history models for each of the *Wollemia* populations identified in the previous steps. *Fastsimcoal2* performs coalescent simulations of the site frequency spectrum (SFS) based on user defined demographic models. The coalescent simulations are performed by sampling from a distribution of potential parameter values, followed by a conditional maximization algorithm (ECM) that optimizes parameters to increase their likelihood given the data ^131^. The DArTSeq dataset for each population was summarized in the form of folded site frequency spectra (i.e. unknown ancestral state of variants, minor SFS), constructed with the *gl2sfs* function in the dartR package ^122^. We implemented a strategy of removing loci and individuals to create a dataset with no missing data, which is a requirement for running fastsimcoal2. After experimenting with different values, we filtered the dataset to SNPs with a maximum of 30% missing data. We then reduced the populations (n samples) to the number of individuals effectively sequenced for the locus with the most missing data (x% missing data) in each population (n*(1-x%)). This was done by randomly sampling n*(1-x%) individuals with no missing data from each locus, resulting in a final dataset of 66 individuals. We transformed the data to tidy format for easier manipulation with the function *tidy_genomic_data* from the radiator package (Gosselin 2020). We then transformed the data back to genind format with the *write_genind* function from the same package. For model testing, we kept one variant per locus to avoid linkage between sites ^132^ and ignored invariant sites (-m option in *fastsimcoal2*). We used the full dataset (invariant sites and secondary variants) to scale parameter values and optimize parameter estimation for the best-fit model identified in the previous step. We manually added the number of invariant sites to the 0 bin in each SFS, which was calculated by subtracting the number of SNPs in the dataset from total number of basepairs (average locus length * number of loci). We ran 50 independent *fastsimcoal2* optimization runs for each demographic model, each run consisting of 40 rounds of ECM with 100,000 coalescent simulations. We tested four single-population models: decrease, increase, bottleneck and constant population size. All parameter search ranges had uniform distributions. The decrease model had four parameters: current size (NCUR, search range from 10-20K), time of population size decrease (TDEC, search range from 10-10K generations), ancestral population size (NANC, 100-100K), and ratio between ancestral and current population (RESIZE). The increase model had four parameters: current population size (NCUR, 1000-100K), time of increase (TINC, 10-10K), ancestral population size (NANC 10-20K) and ratio between ancestral and current population (RESIZE). The bottleneck model had seven parameters: current population size (NCUR, 10-100K), size during bottleneck (NBOT, 1 20K), ancestral size (NANC, 1000-200K), time of bottleneck (TBOT, 10-10K generations), ratio between current and bottleneck population size (RESBOT), ratio between bottleneck and ancestral population size (RESENDBOT) and time of end of bottleneck, considering a standard duration of 100 generations (TENDBOT) ^130^. We selected the model with the lowest Akaike Information Criterion (AIC), calculated after log-transforming the likelihood estimates of the run with the highest likelihood ^131^. We calculated a 95% confidence interval for the parameter estimates of the best model by parametric bootstrapping ^131^. We simulated 100 SFS based on the parameters estimates obtained with the observed SFS. We then performed five optimization runs for each simulated SFS, each consisting of 40 ECM cycles and with 100,000 coalescent simulations.

We also tested joint demographic models for the two populations for which we had the most data (S1 with 28 and S2 with 38 samples, a total of 3600 loci) by using the Approximate Bayesian Computation (ABC) framework associated with Random Forest (RF) algorithms as implemented in DIYABC-RF ^133^. We tested four models: a simple ancestral population split model with no associated population size change (Supplementary Figure 7a); an ancestral population split followed by admixture between the daughter populations (Supplementary Figure 7b); split followed by independent population size decreases in the daughter populations (Supplementary Figure 7c) and a decrease in the ancestral population followed by split into daughter populations (Supplementary Figure 7d). We limited population size changes to decreases in the joint demographic models based on results for the single population models (see **Results**). The priors for all parameters had uniform distributions. Time priors ranged from 100 to 100,000 generations ago, population size priors ranged from 10 to 10,000 individuals, and the admixture prior ranged from 0.01 to 0.5, based on results of admixture analyses. We ran 25,000 and 300,000 simulations per model for the model selection and the posterior estimation steps, respectively, with RFs consisting of 1000 trees. We used default summary statistics to summarize both observed and simulated datasets ^133^. We used 1000 simulated samples for out-of-bag estimation of global and local error measures.

### Genome assembly

We estimated the genome size and heterozygosity of the *Wollemia nobilis*, prior to assembly by counting the 21-mers in the Illumina reads using KMC v3.1.1 ^134^ and analyzing it using a reference-free analysis with GenomeScope v2.0 ^34^. Using the genome size estimate from GenomeScope2, we then selected the longest ∼20x coverage of Nanopore reads with mean qscore greater than QV 10 (phred quality). This was equivalent to selecting reads greater than 40 kbp length (Supplementary Figure 2) for the assembly.

The selected Nanopore sequencing reads were then assembled using the Flye v2.8.1 ^36^ genome assembler, which resulted in 23,120 contigs being assembled with a 9.21 Mbp contig N50 size (11.66Gbp bp of total assembly size). Addressing the consensus accuracy challenges from an ONT assembly, we developed an optimized workflow (Supplementary Figure 2) for polishing the *Wollemia* contigs with a combination of Nanopore reads and short Illumina reads. We performed 2 rounds of polishing with medaka v1.0.3 (-m r941_prom_high_g360) (https://github.com/nanoporetech/medaka) using ONT long reads, 1 round of polishing with POLCA (v3.4.2) ^37^ using short Illumina reads and 2 rounds of Illumina kmer based polishing using our new algorithm HetTrek v0.1 (err_threshold=11, het_threshold=30, unique_threshold=65, anchor_threshold=50, max_nodes_to_search=1000, dist_multiplier=1.2) (https://github.com/kjenike/HetTrek) with kmer size set to k=21, k=51 respectively. Briefly, HetTrek polishing mode removes error sites from assembled contigs by identifying rarely occurring kmers and replacing them with locally assembled sequences. Putative errors are identified based on k-mer counts from raw sequencing reads and a local de Bruijn graph is constructed connecting the non-error sites on either side of the error kmers. The highest coverage path through the de Bruijn graph is used to patch the error site.

### Mitochondrial and chloroplast genome assembly

For the mitochondrial genome, we used a “bait-and-assembly” approach of identifying and assembling those reads that align to the mitochondrial genomes of other closely related species. Specifically, the mitochondrial contigs were assembled using Flye assembler using ONT long reads that are aligned by minimap2 v2.17-r941 ^135^ to other closely related gymnosperm mitochondrial assemblies (^136–139^ and gene set of all gymnosperm mitogenomes from Refseq land plants database. The mitochondrial assembly was polished just as the *Wollemia nobilis* nuclear genome as discussed above. For the chloroplast, we retained the complete *Wollemia nobilis* chloroplast sequence published by ^4,135^ as we were able reproduce their chloroplast sequence by aligning the ONT long reads to their assembly using minimap2 and de-novo assembling them using Flye v2.8.1 assembler.

### Genome quality assessment

We aligned the Nanopore reads to the assembly using minimap2 v2.17-r941 to remove any zero coverage contigs based on nanopore read depth. Furthermore we validated the lack of any bacterial, fungal, viral, protozoan and human contaminants in the final polished assembly using Kraken2 ^140^ metagenomic classifier.

We measured the consensus accuracy and completeness of the final polished assembly via Illumina k-mer copy number spectra using Merqury version 2020-01-29 (^38,140^). Additionally we examined the assembly completeness with conserved single-copy genes using BUSCO v5.0.0 ^141^ using the embryophyta database from OrthoDBv10 ^41^ in the genome mode. Finally we assessed the genome quality using RNASeq mappability to the final polished assembly from 6 different tissue types using the STAR v2.7.5c aligner (--outFilterScoreMinOverLread 0.49 --outFilterMatchNminOverLread 0.49 --outSAMunmapped Within --outSAMprimaryFlag OneBestScore --outSAMstrandField intronMotif --outSAMtype BAM SortedByCoordinate) ^142^.

### Mitochondrial assembly quality assessment

We evaluated the assembled contigs against plant mitochondrial sequences to confirm similarity (BLASTN ^143^ against the “nt” sequence database, best hit approach [e-value < 1e-10]). We then searched the completed assembly for conserved mitochondrial genes, with an emphasis on 41 genes found in other Gymnosperms (*Pinus taeda* (NC_039746.1), *Taxus cuspidata* (MN593023.1), *Cycas taitungensis* (NC_010303.1), *Ginkgo biloba* (NC_027976.1), and *Welwitschia mirabilis* (NC_029130.1)) (TBLASTN ^143^), RefSeq land plant mitochondrial genes against assembly, e-value < 1e-50 & bitscore >=150). Conserved genes absent from the mitochondrial assembly were further searched for in the *Wollemia nobilis* transcriptome and nuclear and chloroplast genome assemblies to check for assembly error.

### Transposable element annotation and insertion times of LTR-RTs

In order to identify transposable elements (TE) across the whole genome, we constructed a de novo TE library for *Wollemia nobilis*.The pipeline extensive *de-novo* TE annotator, EDTA v1.9 ^74^ was ran on five random batches of *Wollemia*’s contigs of about 2Gbp per batch. The pipeline was run without the annotation step with default parameters except for parameter *-curatedlib* where we used the known library of SINE Base v1.1 (https://sines.eimb.ru/) to be able to distinguish SINEs from LTR elements. From each batch we obtained a library of unique elements. We harmonized each TE library using the panTE.pl, a companion script from the EDTA pipeline. The final harmonized TE library was used to further run RepeatMasker v4.1.2-p1 (A.F.A. Smit, R. Hubley & P. Green RepeatMasker at http://repeatmasker.org) using the parameters recommended by the EDTA author against each batch to recover any of the TEs that are split between two contigs. We performed a final EDTA run from the annotation step using all files created before including the parameter *-rmout* using the *.out file from the RepeatMasker and the parameter *--overwrite 0.* This last step served to annotate TEs and calculate TE abundance for each batch.

In order to estimate LTR-RT insertion times we used the result file *mod.pass.list containing the intact LTR-RTs merged in one file from all the genome batches. Each file is a result from the LTR-RETRIEVER pipeline within the EDTA pipeline per genome batch. This file was reanalyzed using a Python script, that recomputes the insertion times (*T*) using the formula *T=K/(2*r)*. The mutation rate (*r*) was set at 2.2 × 10^-9^ per site per year ^39^. K was computed as 1-identity%, taken from the “Identity” column in *mod.pass.list file.

### Gene annotation and gene set assessment

Raw RNASeq read quality was assessed using FastQC v0.11.9 (https://www.bioinformatics.babraham.ac.uk/projects/fastqc/), reference based transcripts were generated using the STAR-Stringtie2 v2.1.2 ^144^ workflow and de novo transcriptome assembly was constructed using Trinity(V) ^145^ transcriptome assembler. Furthermore, we filtered out the invalid splice junctions generated by STAR aligner using portcullis v1.2.0 ^144,146^ method. Additionally we also lifted orthologs from several other published gymnosperm genomes using the Liftoff v1.6.1(-exclude_partial-copies) ^147^ pipeline. These results along with transcripts mapped to genome using gmap version 2020-10-14 ^148^ from several closely related species from treegenesdb.org were used as EST evidence in addition to the protein evidence from the uniprot/swissprot database and refseq spermatophyta databases. Structural gene annotations were then generated using the Mikado v2.0rc2 ^148,149^ framework using the evidence set provided following the Snakemake-based pipeline, Daijin. Functional annotation of the mikado gene models was performed using the ENTAP v0.10.8 ^49^ pipeline using the eggNOG5.0 ^150^, Uniprot/Swissprot ^151^, TREMBL, RefSeq all/plants and Spermatophyta proteins database ^151,152^, InterProScan5 ^50^ using Pfam, TIGRFAM, Gene ontology options and TRAPID ^51^.

We assessed the completeness of the gene models by benchmarking single copy orthologs using BUSCO5 in protein mode against embryophyta_odb10, tracking the placement of RNASeq reads with the genic regions across all the tissue types using STAR and qualimap v.2.2.1 ^49,153^ and verifying the presence of custom selected gymnosperm specific orthologs within the annotated gene models.

The NLR-Annotator pipeline ^154^ was deployed to identify potential NBS-LRR genes.

### Gene family evolution analyses

We analyzed the predicted coding sequences from the genome assemblies of *Wollemia nobilis* and seven other species which cover the major clades of land plants. Our phylogenomic pipeline PhyloGeneious (github.com/coruzzilab/PhyloGeneious) was used to infer groups of orthologs, gene families, and one-to-one orthologs^155^. Our taxon set include gymnosperms (*Sequoiadendron giganteum* (TreeGenes v2.0), *Pinus tabuliformis* (CNGB 2021), *Ginkgo biloba* (CNGB 2021)), angiosperms (*Amborella trichopoda* (Plaza v1.0), *Arabidopsis thaliana* (TAIR Araport11)), lycophytes (*Selaginella moellendorffii* (Phytozome v1.0)), and non-vascular plants (*Physcomitrella patens* (Phytozome v3.0)). Amino acid sequences of select gene families were aligned in M-Coffee ^156^ by combining the output of T-Coffee, MAFFT, and MUSCLE. Phylogenetic relationships among homologs were estimated in IQ-TREE v2.1.2 ^157^ with internode branch support obtained using the ultrafast bootstrap approximation and 5000 replicates ^158^, and RAxML v8.2.12 with branch support estimated with 500 rapid bootstrap replicates ^159^. Families of orthologs exceeding 1000 members were analyzed using FastTree v2.1.11 ^160^ using the LG+Γ substitution model and the Shimodaira-Hasegawa-like local node support implementation ^161^.

To identify the *APETALA2*, *AINTEGUMENTA, RAV and ERF* homologs in *W. nobilis*, we performed a BLAST search of the genome using as query homologs previously reported across land plants (Kim et al., 2006; Zumajo-Cardona and Pabón-Mora, 2016). A total of 344 protein sequences were aligned in MAFFT ^162^. According to jModelTest2 ^163^, GTRGAMMA was identified as the best-fit substitution model. RAxML v8.0.0 was used to estimate phylogenetic relationships ^124^. Six closely related genes belonging to the ARF2 were used as outgroup: (ARF2: NP_851244.1; *Arabidopsis lyrata* (ArlyARF2): AL8G38770.t1; *Arabidopsis helleri* (ArhaARF2): Araha.4168s0011.1; *Boechera stricta* (BostARF2): Bostr.26833s0038.1; *Capsella rubella* (CaruARF2): Carubv10025860m; *Capsella grandiflora* (CaruARF2): Cagra.4553s0009.1) were used as the outgroup. Two separate analyses were performed for the two subclades, AP2/ANT lineage with a total of 186 sequences, and the ERF lineage with a total of 146 sequences.

## Acknowledgements

This work was funded by the National Science Foundation Plant Genome Research Program IOS grant 1758800 to R.W.M., R.A.M., G.M.C., K.V., M.C.S., B.A., D.P.L., S.-O.K. We also acknowledge, the National Institutes of Health (U24-HG006620) and Howard Hughes Medical Institute to R.M., NSF PGRP Grant IOS 0922738 and the Zegar Family Foundation Grant A16-0051 to G.M.C., fellowship funding to V.M.S from NIH1T32 GM132037, NSF OPUS Grant NSF-DEB 2140319 to D.W.S. The authors acknowledge assistance from the Cold Spring Harbor Laboratory Shared Resources, funded by NIH (grant S10OD028632-01). S.N. and S.M. received funding from the European Union’s Horizon 2020 research and innovation programme under the Marie Skłodowska-Curie grant agreement no. 101007738. W.R.M is the Davis Family Professor of Human Genetics at CSHL. G.M.C is the Carroll & Milton Professor of Biology at NYU. D.W.S. is the Cullman Senior Curator Emeritus at NYBG and President of the Torrey Botanical Society. We thank Barbara Baldan for her help with collecting samples at the University of Padova. We thank the Royal Botanic Gardens and Domain Trust of Australia, for funding the genome resequencing, along with Illumina Inc. that contributed some consumables. We thank Newcastle Museum, NSW, Australia, for access to relevant fossil specimens, and Yin Peng Lee and Stella Loke for sample processing and sequencing of the Wollemi pine population in Australia.

## Author contributions

W.R.M., R.A.M., L.J.C., G.M.C., D.W.S., K.V., S-O.K., M.C.S, B.A., D.P.L. designed the study. B.A., S.F. and D.W.S collected the Wollemia samples at NYBG and S.N. and S.M collected Wollemia samples at University of Padova. H.M., M.R., L.J.C collected Wollemia population samples in Australia. S.G., S.F., C.S.A. processed samples and generated the data. S.R., K.J., and M.K. performed the genome assembly. S.R., K.V, S-O.K, G.E., V.M.S, M.S.K., B.A., C.Z-C. C.S.A. performed transcriptomics annotation and analysis. O.M.R, S.O., S.R., and C.S.A. annotated and analyzed TEs. C.S.A. generated data annotated and analyzed sRNA. C.S.A. generated and analyzed the methylome data. L.J.C., H.M., M.R generated the population data. L.A.C., S-O. K performed the population analysis. All authors contributed with ideas and discussion, and drafting of the manuscript.

## Data availability

The Genome data has been deposited in GEO under accession number PRJNA564454.

