## Supplemental Figures for "The genome of the Wollemi pine, a critically endangered “living fossil” unchanged since the Cretaceous, reveals extensive ancient transposon activity"

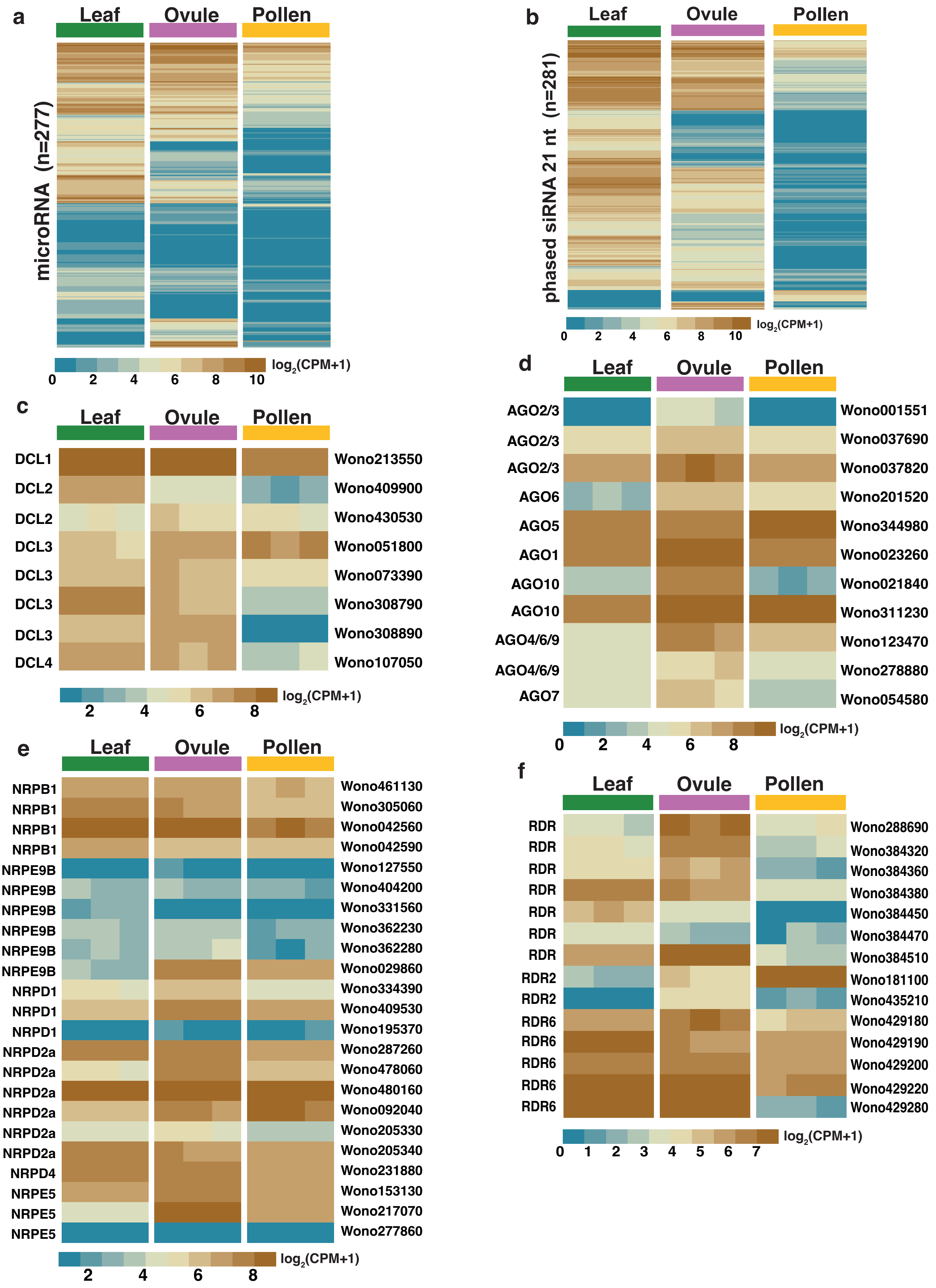


**Extentded Data Figure 1 l miRNA, 21-nt phasiRNA and related putative enzyme expression in leaf, ovule, and mature pollen grains.** **a,** Expression of all annotated miRNA. **b,** Expression of all annotated 21-nt phased siRNAs. **c**, Expression of the putative *Dicer-like* (*DCL*) genes. **d**, Expression of the putative *Argonaute* (*AGO*) genes. **e**, Expression of the putative RNA polymerases subunits **f**, Expression of the putative *RNA-dependent RNA polymerases* (RDR) genes.

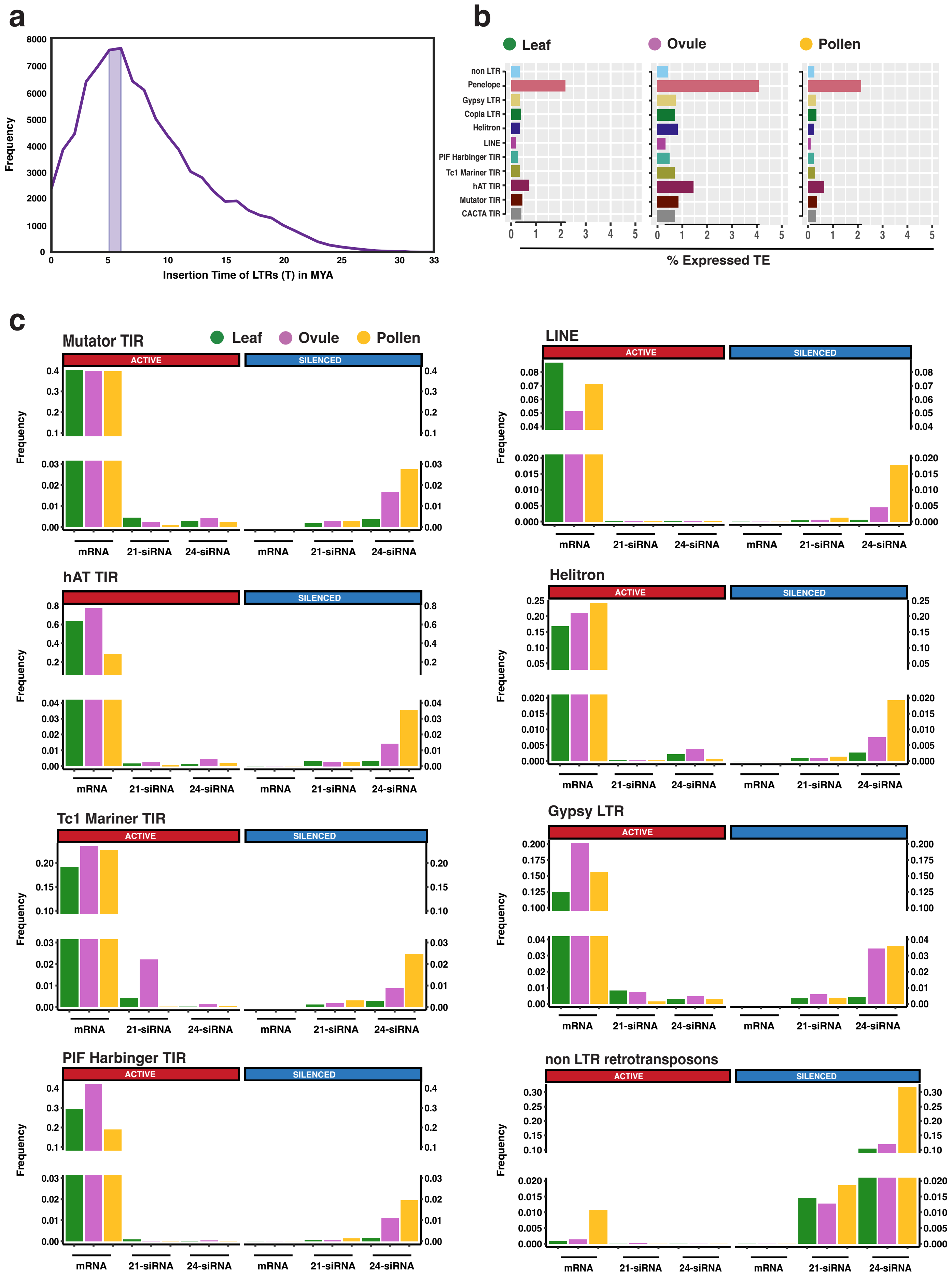


**Extended Data Figure 2 l Transposable elements expression in leaf, ovule, and mature pollen grains. a,** LTR Insertion time. **b**, Percentage of active TEs. **c**, TEs classes mRNA, 21 and 24-nt TE-derived siRNAs expression frequency (sum of CPM / number of TEs).

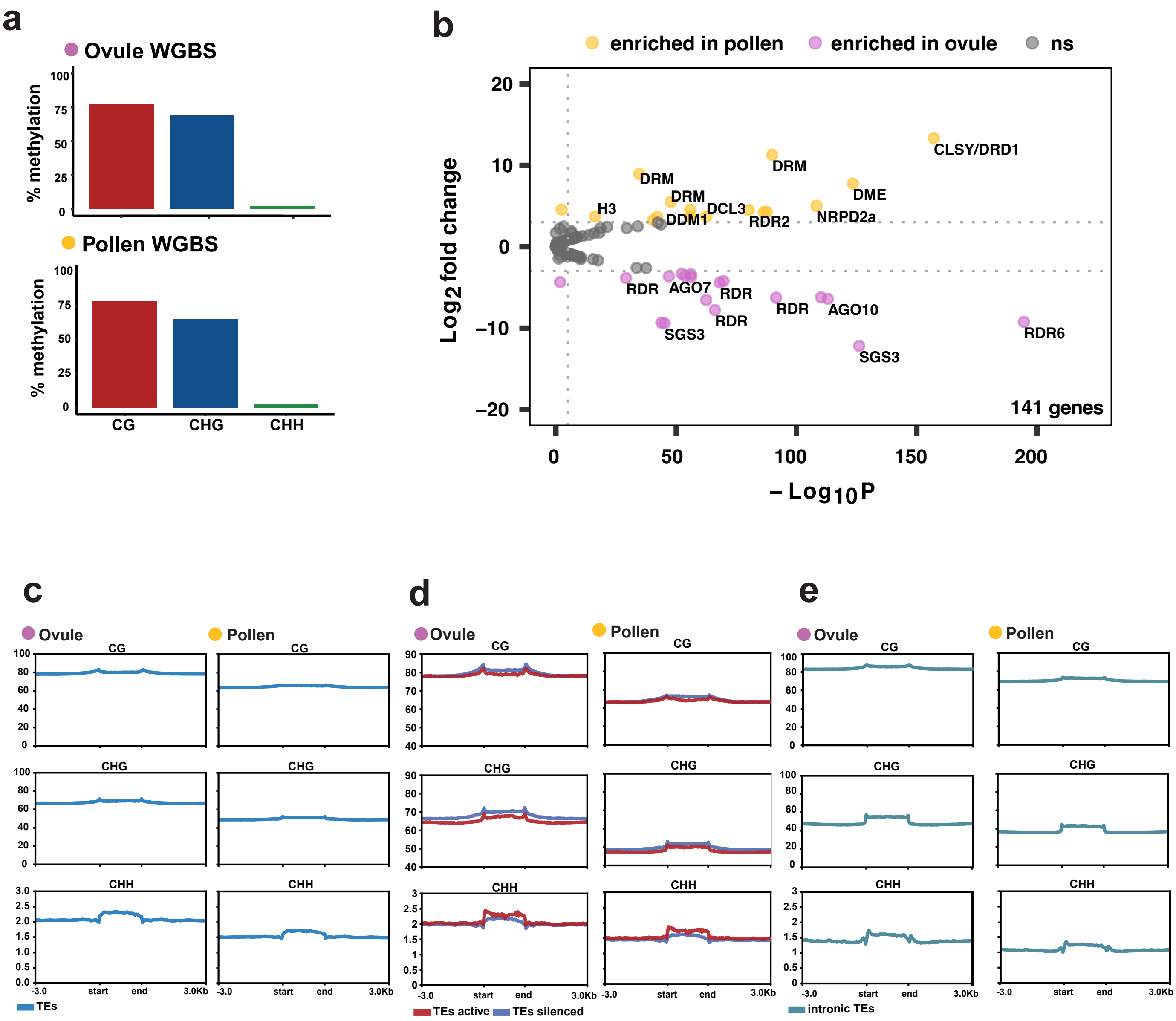


**Extended Data Figure 3 I DNA methylation in ovule, and mature pollen grains. a**, Total percentage of DNA CG, CHG, and CHH methylation. **b**, Epigenetic genes up regulated in both tissues. **c**, DNA methylation in all TEs. **d**, DNA methylation in all silenced (blue) and active (red) TEs. **e,** DNA methylation in all intronic TEs.

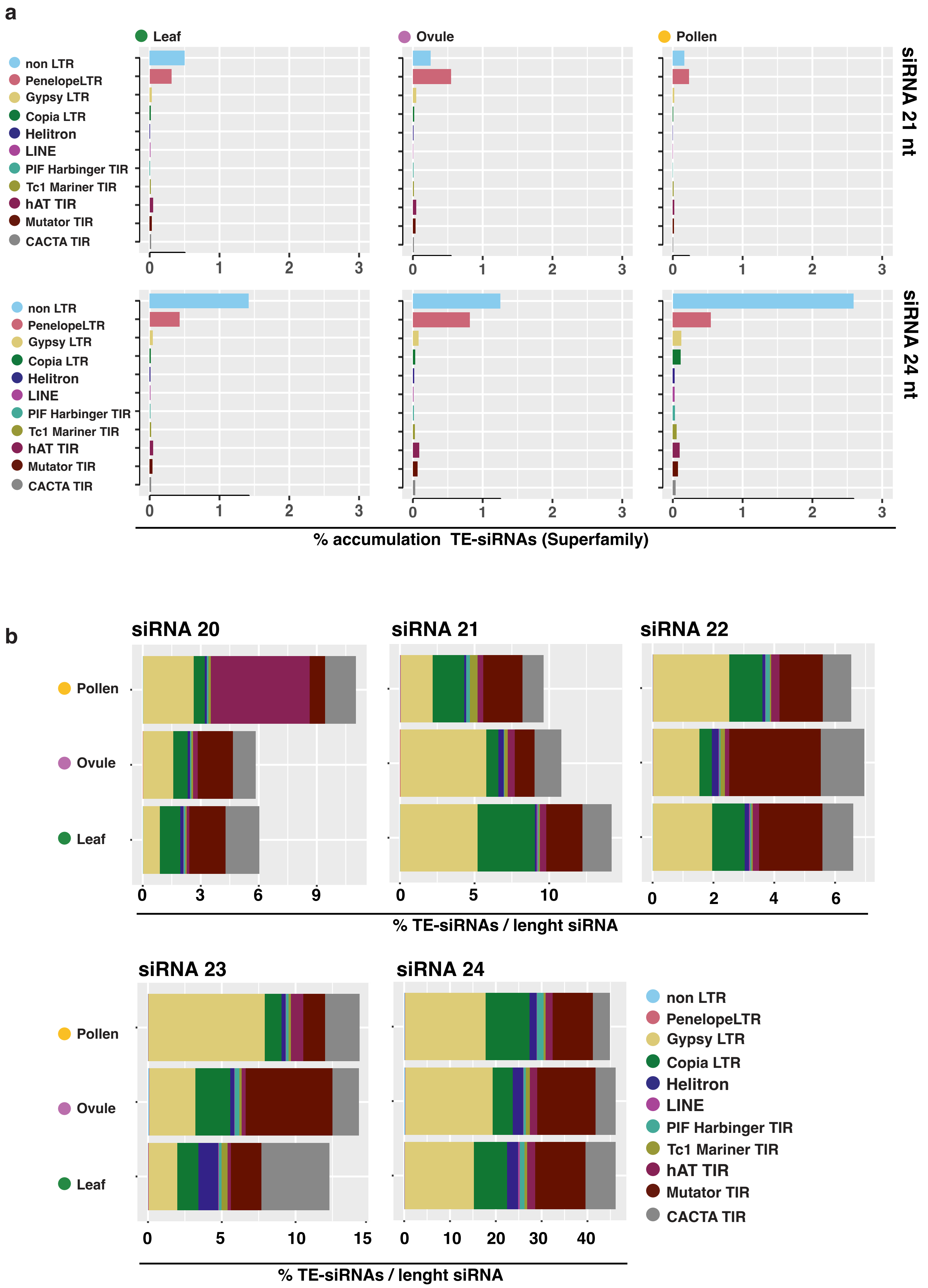


**Extended Data Figure 4 l TE derived siRNAs accumulation in leaf, ovule, and mature pollen grains. a**, Percentage of elements generating 21nt and 24nt-siRNA. **b**, Percentage of 20 to 24nt-siRNAs derived fromTEs.

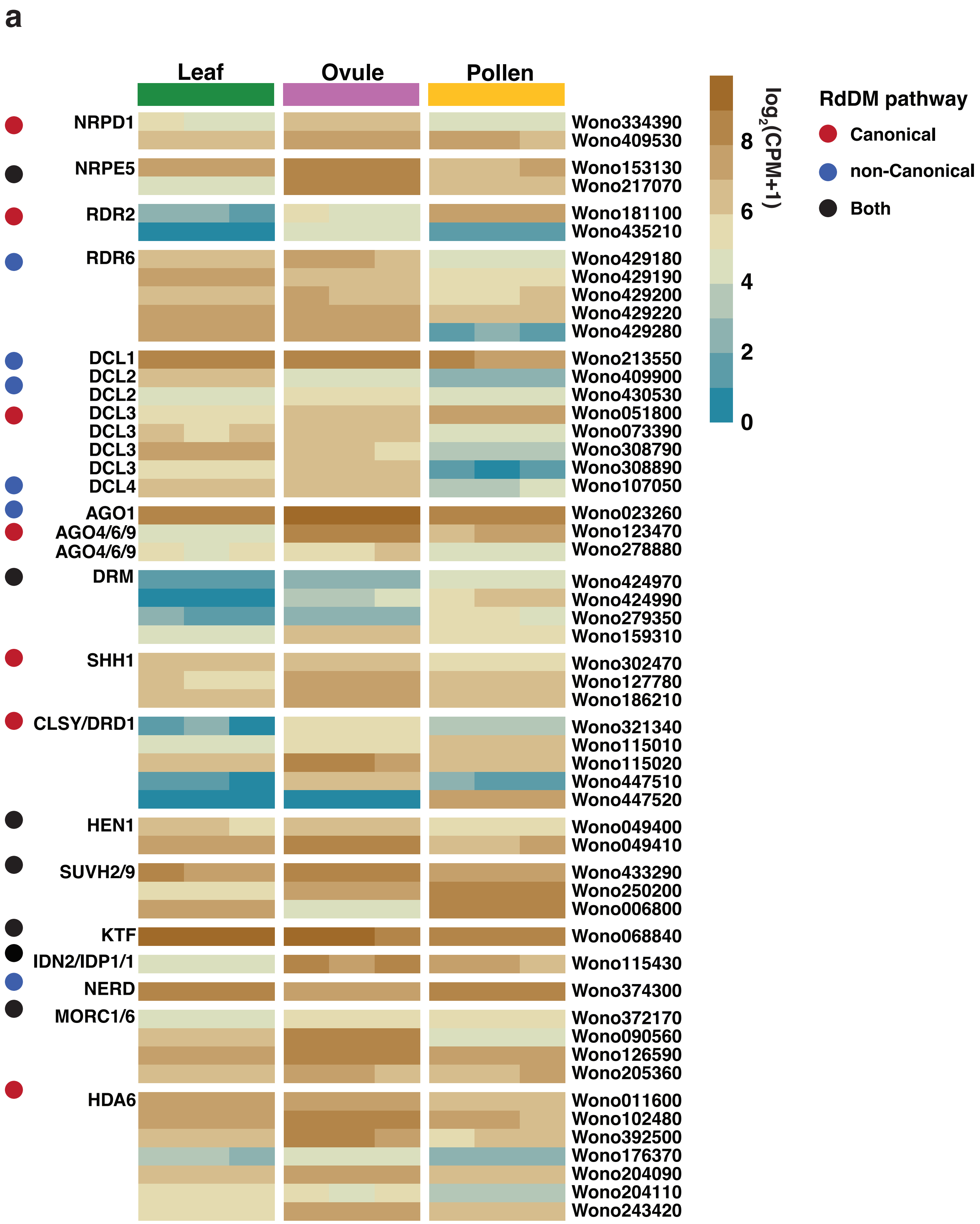


**Extended Data Figure 5 I Active RdDM pathway in leaf, ovule, and mature pollen grains. a**, Putative homologs RdDM genes expression, divided into canonical (red dot), non-canonical (blue dot), and both (black dot) pathways.

a

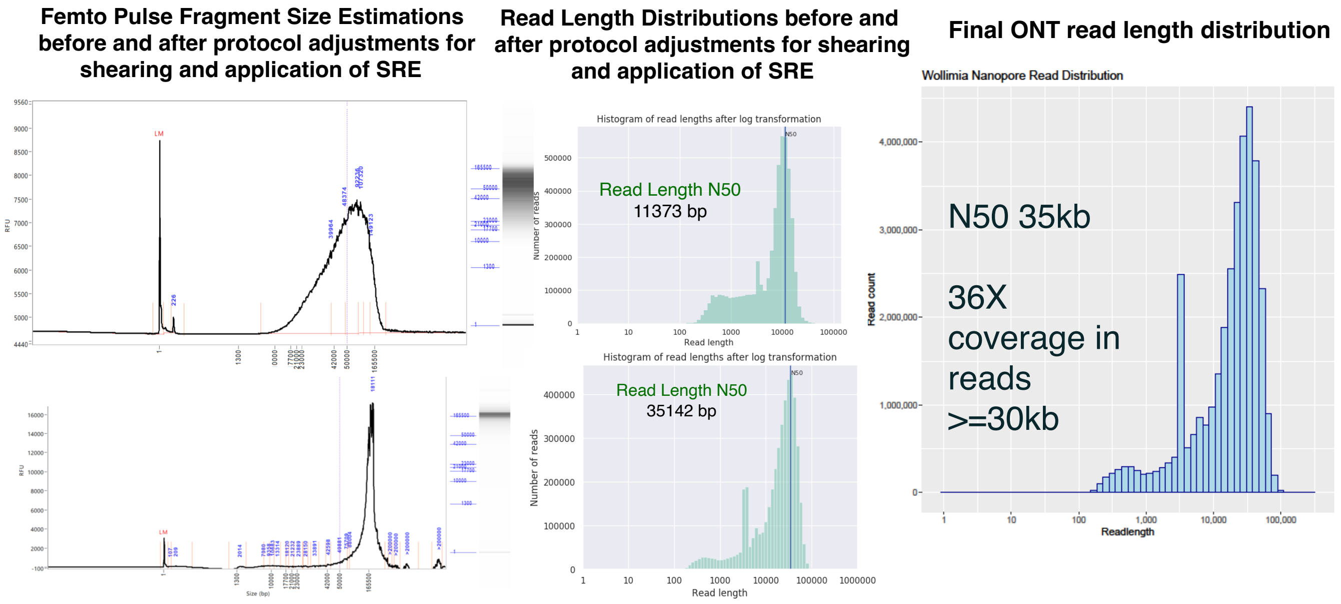


b

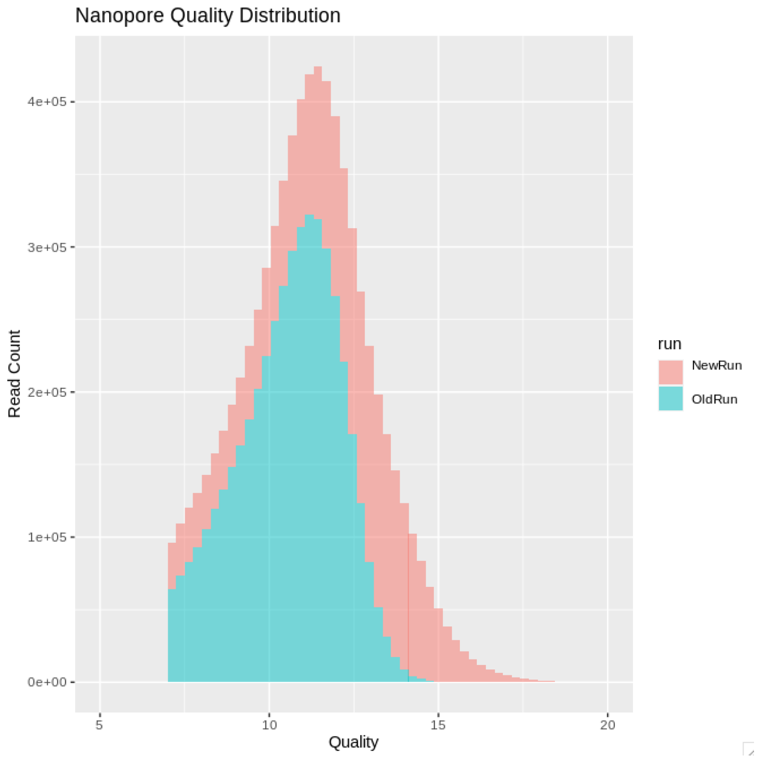


**Supplementary Figure 1 I Assembly Pipeline, Genomescope and ONT Read Coverage. a,** Optimization of DNA fragment size selection for targeting long (>=30kb) reads for ONT sequencing. Improved shearing parameters and use of the Short Read Eliminator kit increased the proportion of long reads as indicated by the Femto Pulse traces and sequenced read length histograms. **b,** Histogram of read quality improvements after re-processing. Re-basecalling with the new version of GuppyV4 improved raw read accuracy over the original basecalls (GuppyV3).

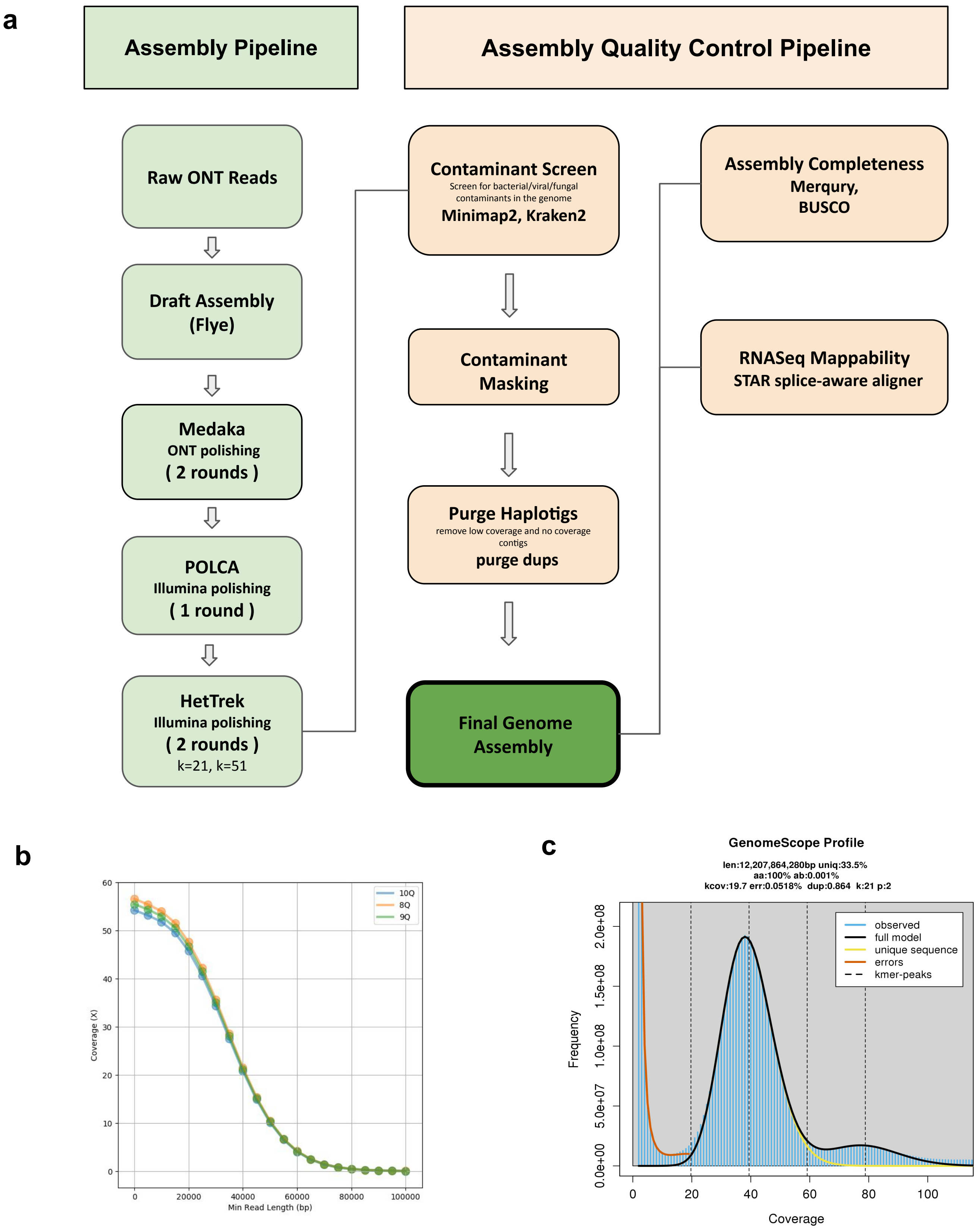


**Supplementary Figure 2 I Assembly Pipeline, Genomescope and ONT Read Coverage. a,** Flowchart showing the assembly processing steps in generating *W.nobilis* 1.0. **b,** Coverage distribution of ONT reads at varying lengths and read quality values 9Q (green),8Q (red),10Q (blue). **c,** K-mer spectra and model fitted by Genomescope 2.0 based based on the short Illumina reads of *W.nobilis*.

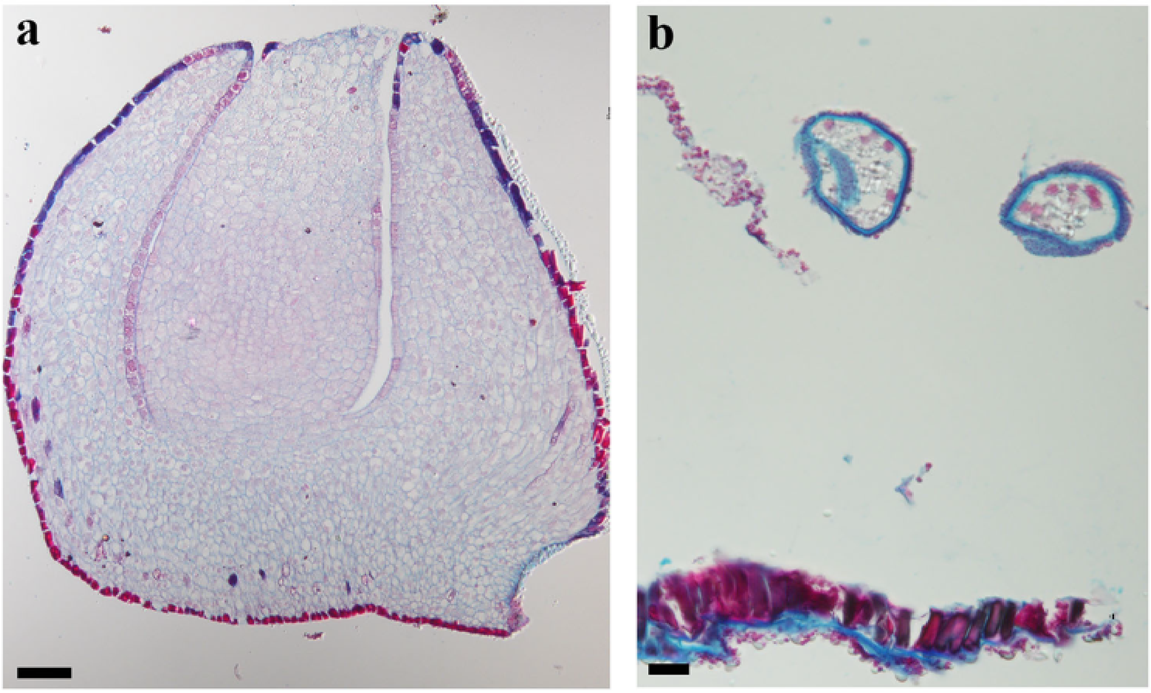


**Supplementary Figure 3 l Ovule and pollen cone anatomy.** Safranin red and astra blue staining of sections of ovules  and pollen cones used for RNA transcriptomes**,** small RNA and methylome analyses, showing anatomy and developmental stages of **a,** ovules (scale bar = 100 $\mu$m) and **b,** pollen (scale bar = 50 $\mu$m).

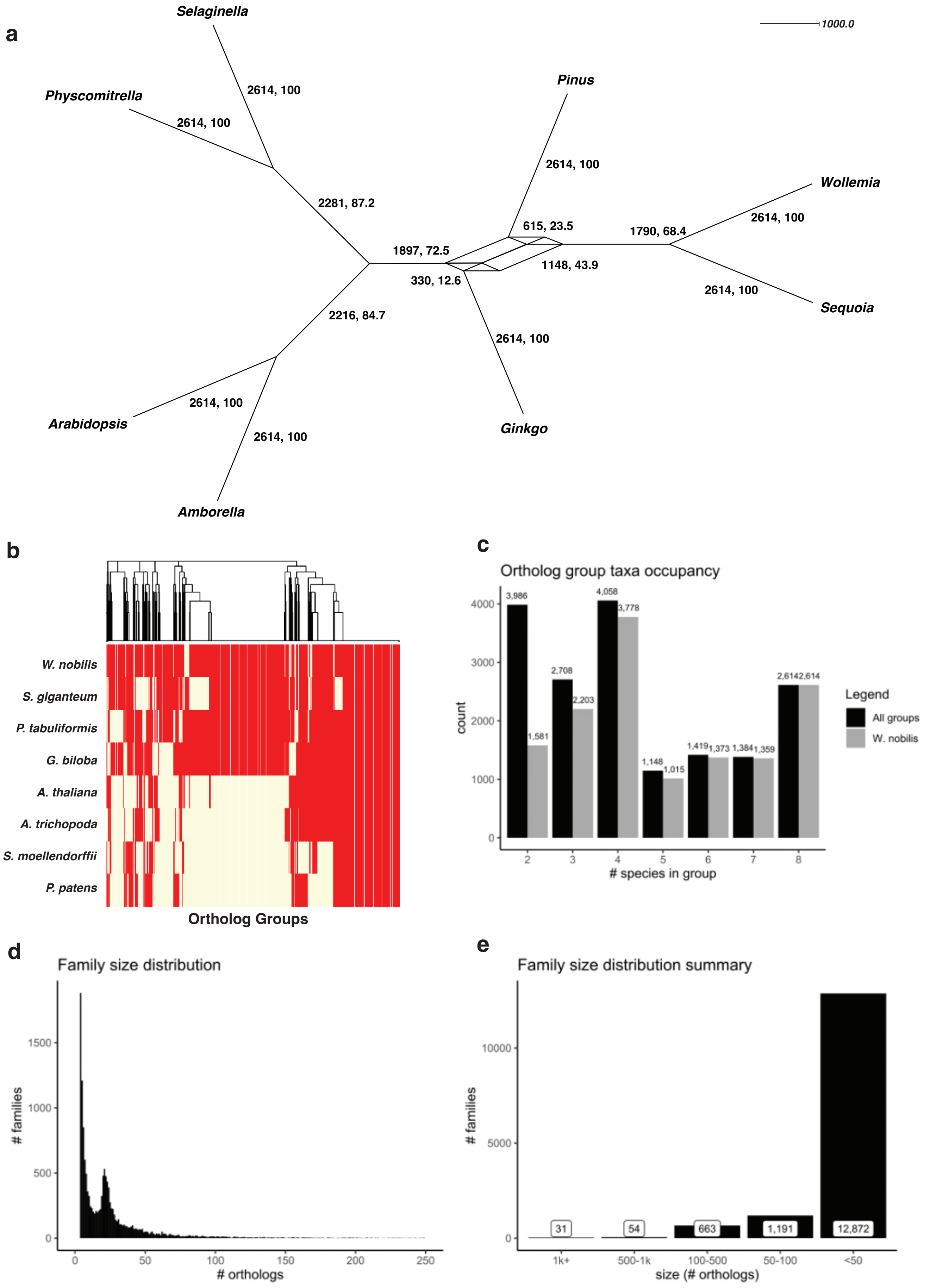


**Supplementary Figure 4 I Shared gene and gene family distributions across land plants. a,** Consensus network of 2,614 phylogenetic trees from the 2,614 1:1 orthologs shared across embryophytes. **b,**Presence/absence of orthologous genes found in three or more species. Tracheophytes (i.e. all but *Physcomitrium patens*), spermatophytes, and gymnosperms had 232, 725, and 3,331 shared groups of orthologs, respectively. **c,** Distribution of ortholog groups based on the number of species represented. Gray bars indicate the number of groups containing orthologs from *Wollemia nobilis*. A large number of conserved, gymnosperm-specific ortholog groups is shown. **d–e,** Distribution of 10,210 family clusters by the number of orthologs. The largest gene family contained 26,093 orthologs, yet the majority of families contain less than 50 orthologs.

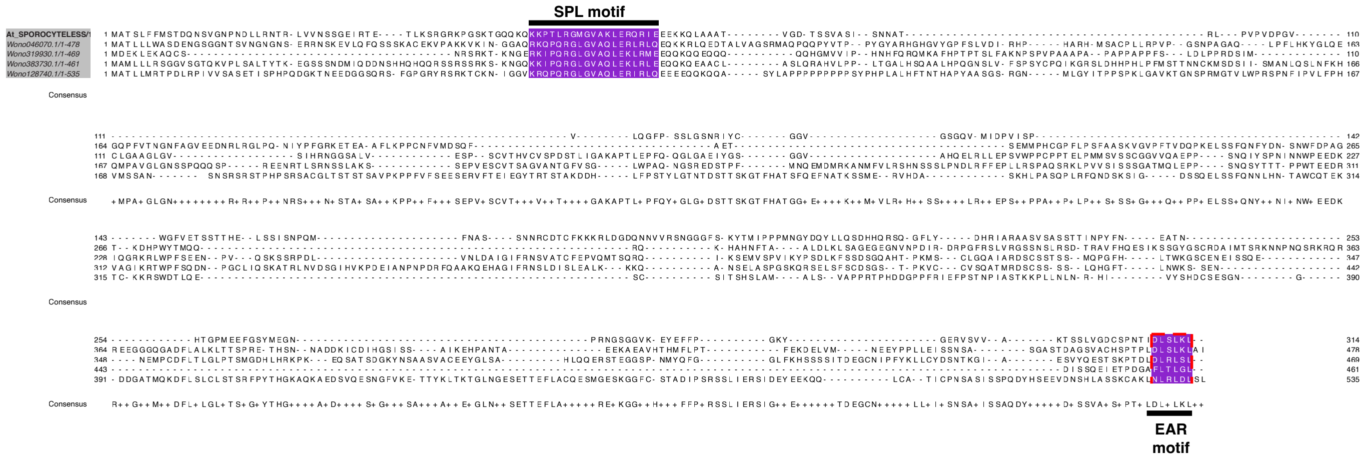


**Supplementary Figure 5 | *Wollemi nobilis* genome encodes mutliple putative orthologs of SPOROCYTE-LESS/NOZZLE transcription factor**. Homology search of the predicted proteome for *Wollemi nobilis* identified four gene models that encode a SPL/NZL transcription factor. All four gene models include the 5’ SPL motif as well as the 3’ EAR motif.

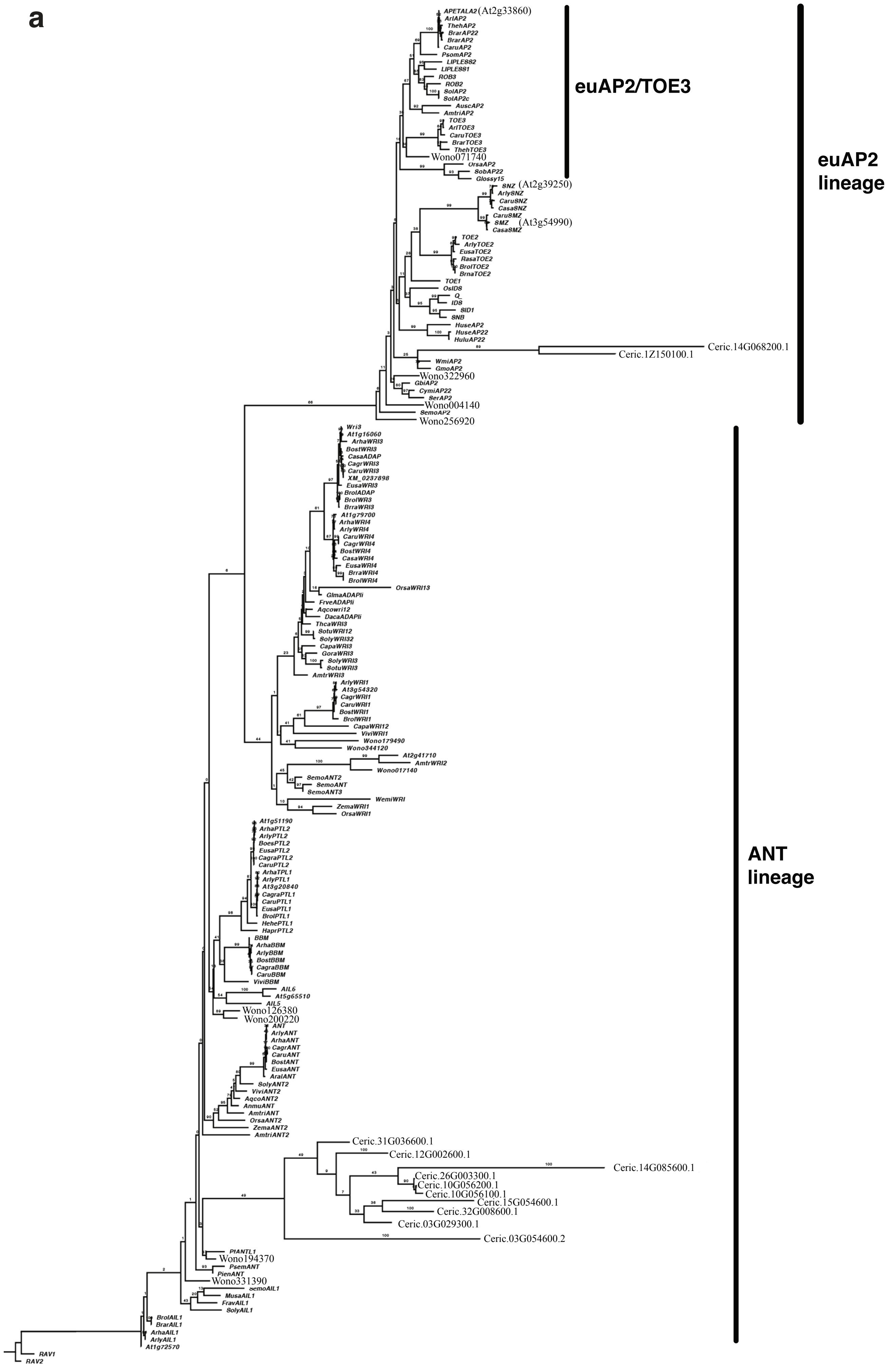
a
euAP2/TOE3
euAP2
lineage
ANT
lineage
APETALA2(At2g33860)
Wono071740
SNZ (At2g39250)
SMZ (At3g54990)
Ceric.14G068200.1
Ceric.1Z150100.1
Wono322960
Wono004140
Wono256920
Wono179490
Wono344120
Wono017140
Wono126380
Wono200220
Ceric.31G036600.1
Ceric.12G002600.1
Ceric.14G085600.1
Ceric.26G003300.1
Ceric.10G056200.1
Ceric.10G056100.1
Ceric.15G054600.1
Ceric.32G008600.1
Ceric.03G029300.1
Ceric.03G054600.2
Wono194370
Wono331390
RAV1
RAV2

**b**

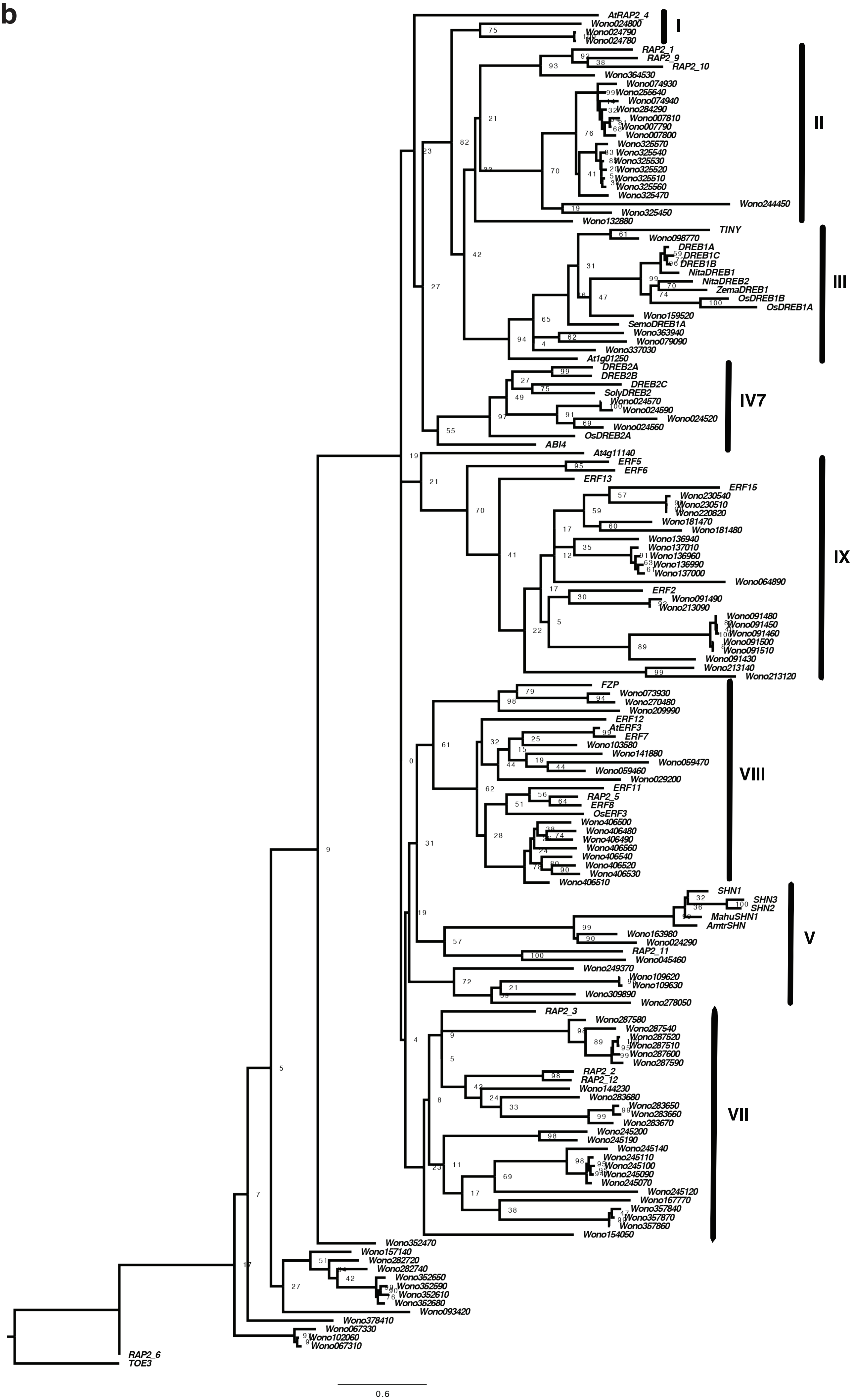
AtRAP2_4
Wono024800
Wono024790
Wono024780
I
RAP2_1
RAP2_9
RAP2_10
Wono364530
Wono074930
Wono255640
Wono074940
Wono284290
Wono007810
Wono007790
Wono007800
Wono325570
Wono325540
Wono325530
Wono325520
Wono325510
Wono325560
Wono325470
Wono244450
Wono325450
Wono132880
II
TINY
Wono098770
DREB1A
DREB1C
DREB1B
NitaDREB1
NitaDREB2
ZemaDREB1
OsDREB1B
OsDREB1A
Wono159520
SemoDREB1A
Wono363940
Wono079090
Wono337030
At1g01250
III
DREB2A
DREB2B
DREB2C
SolyDREB2
Wono024570
Wono024590
Wono024520
Wono024560
OsDREB2A
ABI4
IV7
At4g11140
ERF5
ERF6
ERF13
ERF15
Wono230540
Wono230510
Wono220820
Wono181470
Wono181480
Wono136940
Wono137010
Wono136960
Wono136990
Wono137000
Wono064890
ERF2
Wono091490
Wono213090
Wono091480
Wono091450
Wono091460
Wono091500
Wono091510
Wono091430
Wono213140
Wono213120
IX
FZP
Wono073930
Wono270480
Wono209990
ERF12
AtERF3
ERF7
Wono103580
Wono141880
Wono059470
Wono059460
Wono029200
ERF11
RAP2_5
ERF8
OsERF3
Wono406500
Wono406480
Wono406490
Wono406560
Wono406540
Wono406520
Wono406530
Wono406510
VIII
SHN1
SHN3
SHN2
MahuSHN1
AmtrSHN
Wono163980
Wono024290
RAP2_11
Wono045460
Wono249370
Wono109620
Wono109630
Wono309890
Wono278050
V
RAP2_3
Wono287580
Wono287540
Wono287520
Wono287510
Wono287600
Wono287590
RAP2_2
RAP2_12
Wono144230
Wono283680
Wono283650
Wono283660
Wono283670
Wono245200
Wono245190
Wono245140
Wono245110
Wono245100
Wono245090
Wono245070
Wono245120
Wono167770
Wono357840
Wono357870
Wono357860
Wono154050
VII
Wono352470
Wono157140
Wono282720
Wono282740
Wono352650
Wono352590
Wono352610
Wono352680
Wono093420
Wono378410
Wono067330
Wono102060
Wono067310
RAP2_6
TOE3
0.6

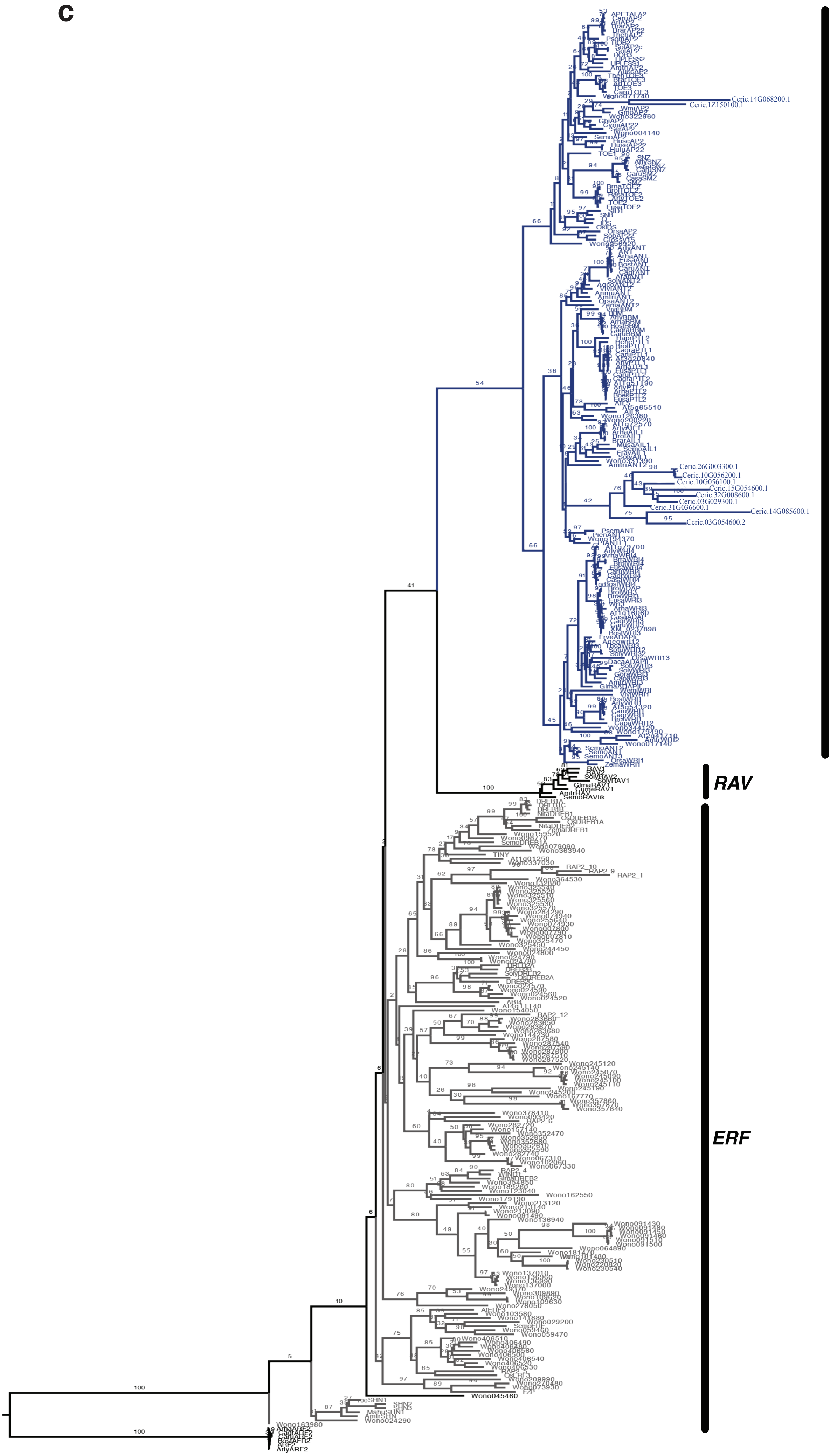
c
AP2/ ANT
RAV
ERF

**d**

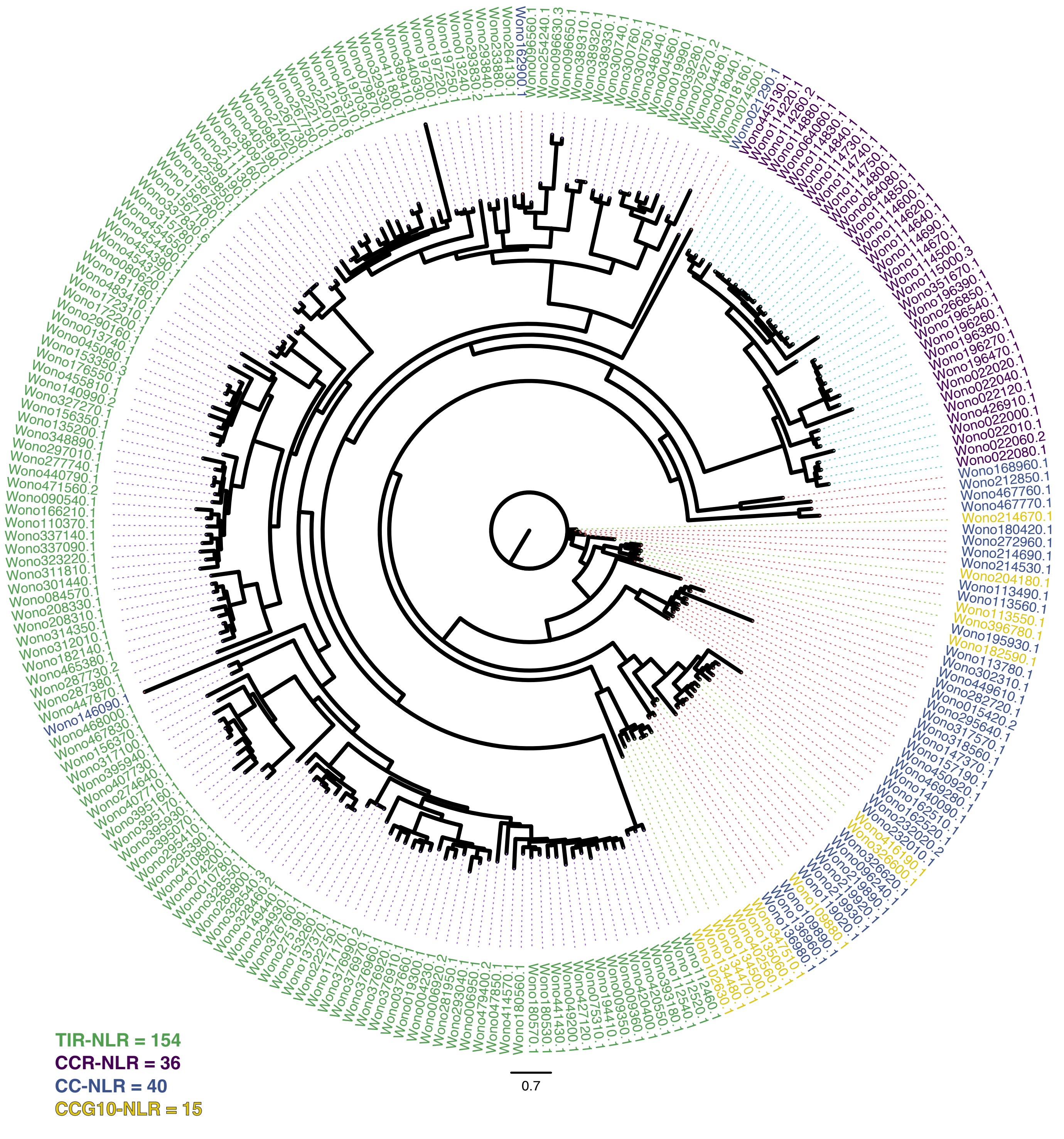

e

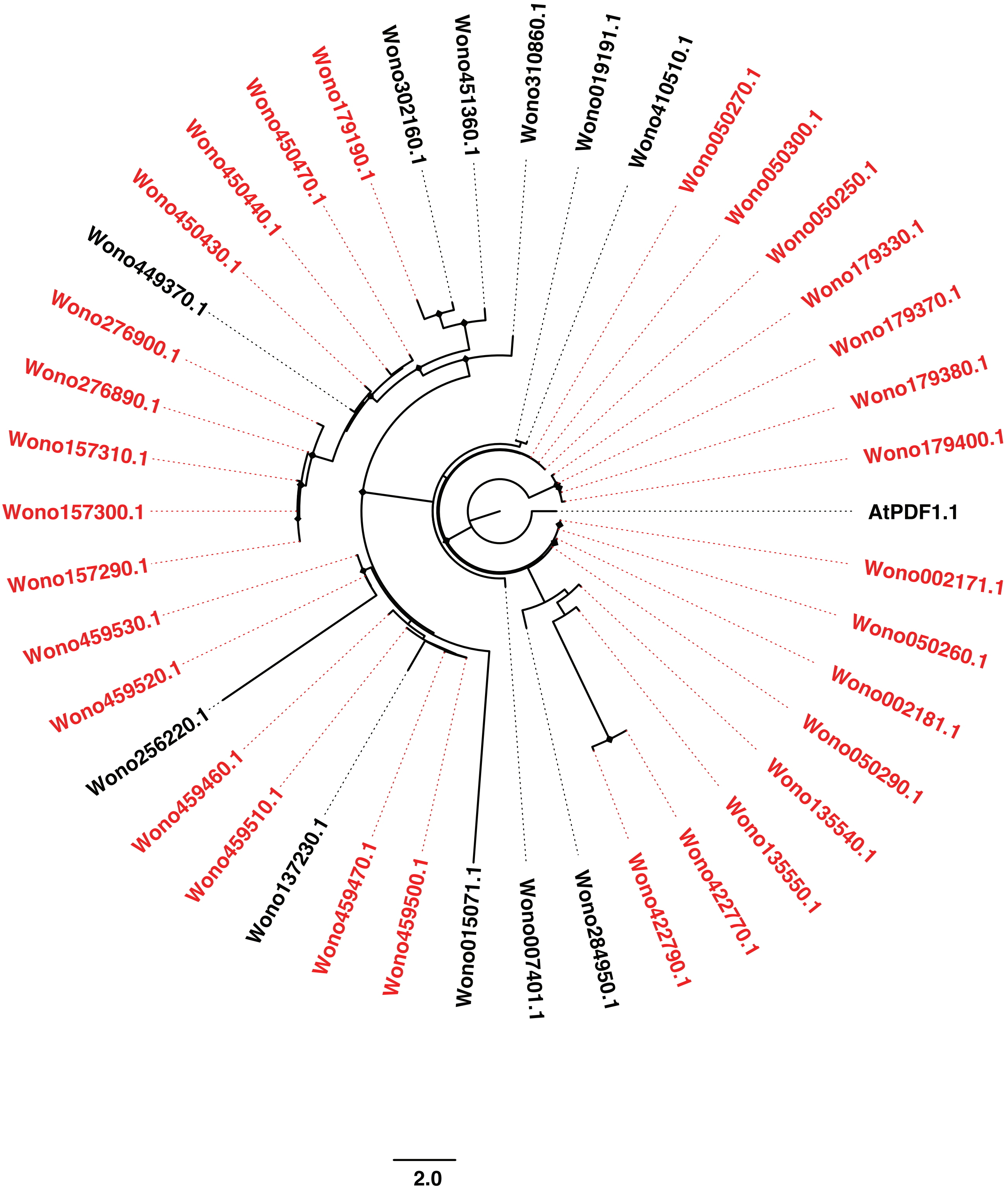
Wono302160.1
Wono451360.1
Wono310860.1
Wono019191.1
Wono410510.1
Wono050270.1
Wono050300.1
Wono050250.1
Wono179330.1
Wono179370.1
Wono179380.1
Wono179400.1
AtPDF1.1
Wono002171.1
Wono050260.1
Wono002181.1
Wono050290.1
Wono135540.1
Wono135550.1
Wono422770.1
Wono422790.1
Wono284950.1
Wono007401.1
Wono015071.1
Wono459500.1
Wono459470.1
Wono137230.1
Wono459510.1
Wono459460.1
Wono256220.1
Wono459520.1
Wono459530.1
Wono157290.1
Wono157300.1
Wono157310.1
Wono276890.1
Wono276900.1
Wono449370.1
Wono450430.1
Wono450440.1
Wono450470.1
Wono179190.1
2.0

f

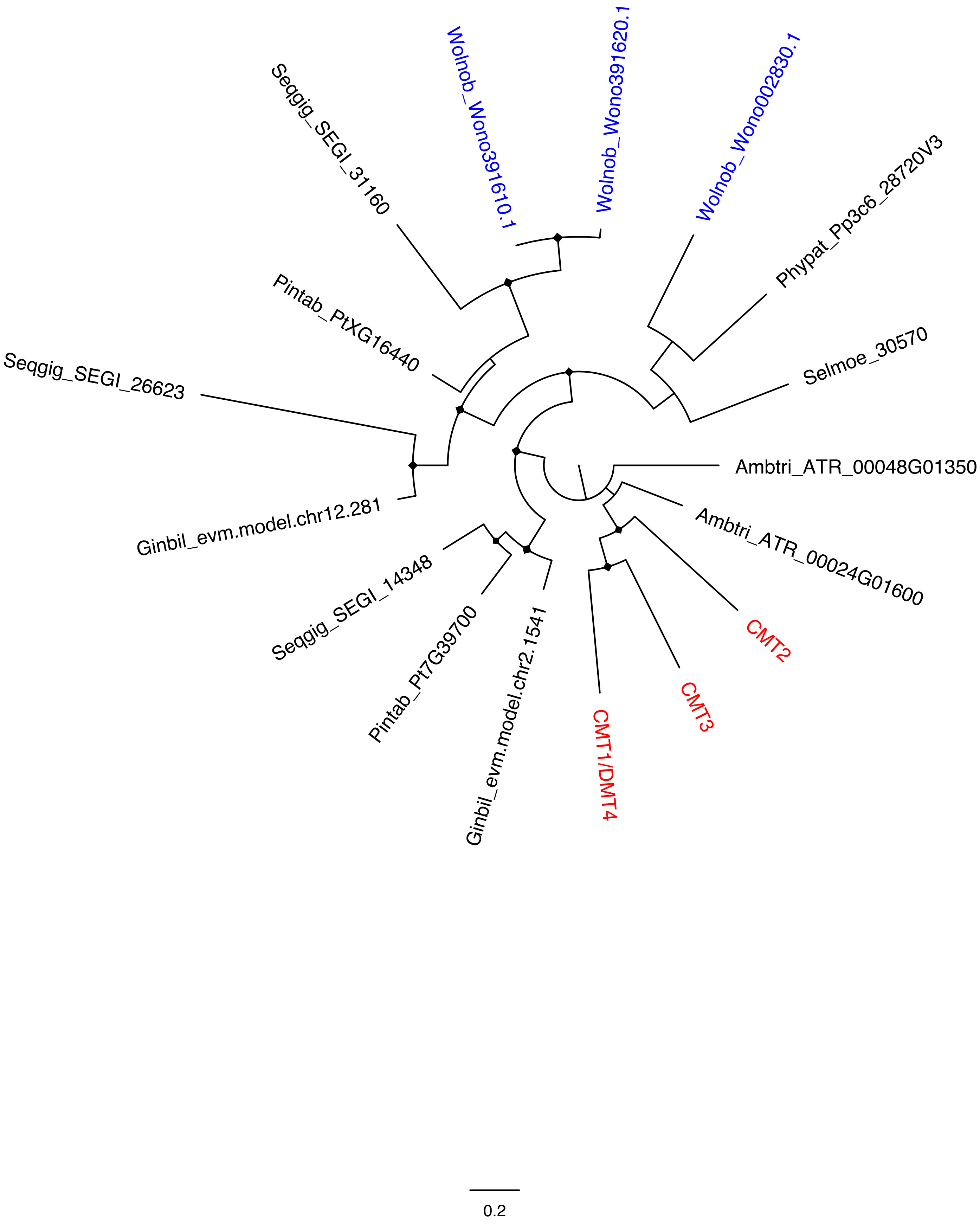
Wolnob_Wono391610.1
Wolnob_Wono391620.1
Wolnob_Wono002830.1
Seqgig_SEGI_31160
Phypat_Pp3c6_28720V3
Pintab_PtXG16440
Selmoe_30570
Seqgig_SEGI_26623
Ambtri_ATR_00048G01350
Ginbil_evm.model.chr12.281
Ambtri_ATR_00024G01600
Seqgig_SEGI_14348
CMT2
Pintab_Pt7G39700
CMT3
Ginbil_evm.model.chr2.1541
CMT1/DMT4
0.2

**g**

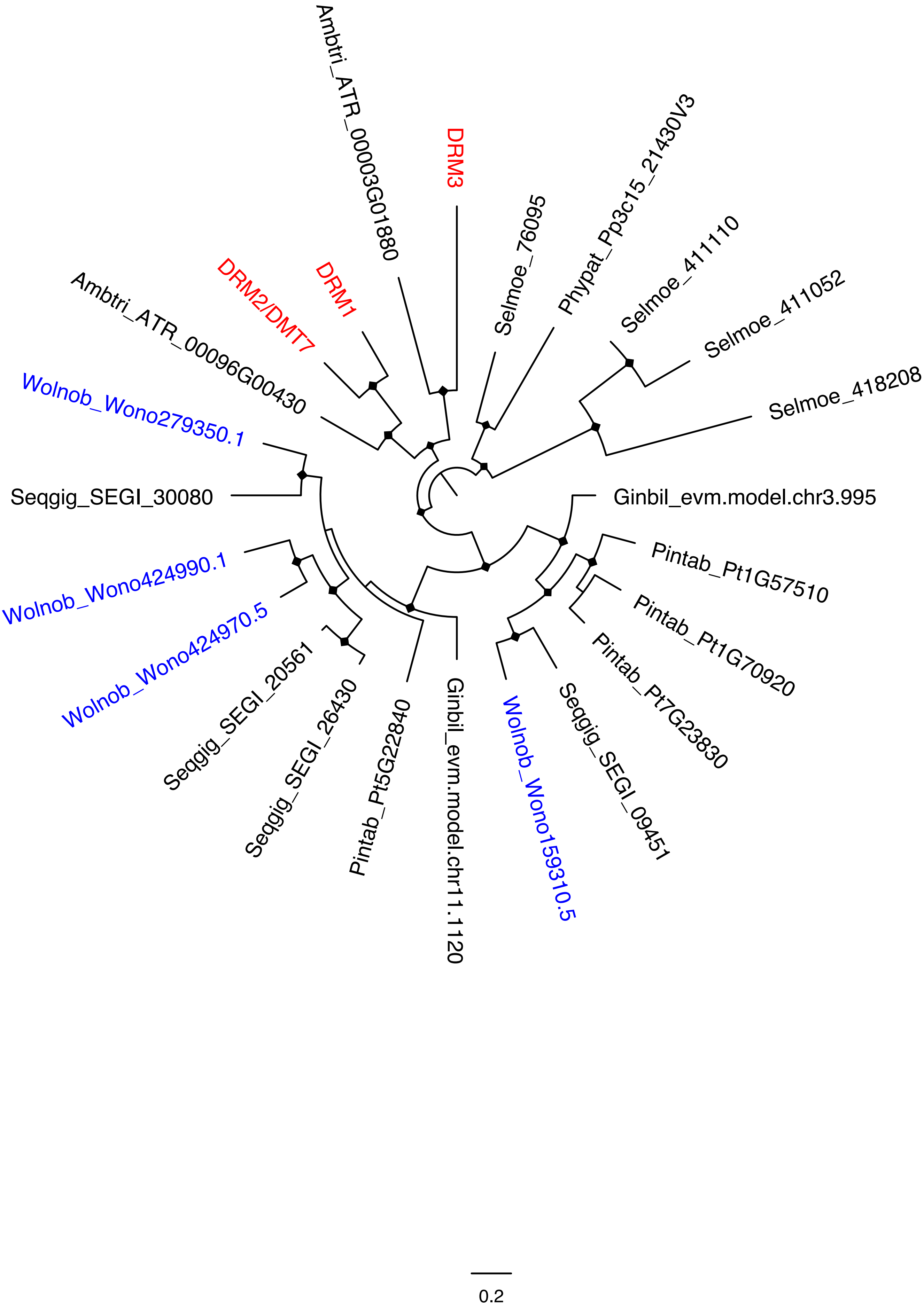

Ambtri_ATR_00003G01880
DRM3
Selmoe_76095
Phypat_Pp3c15_21430V3
Selmoe_411110
Selmoe_411052
Selmoe_418208
DRM1
DRM2/DMT7
Ambtri_ATR_00096G00430
Wolnob_Wono279350.1
Seqgig_SEGI_30080
Ginbil_evm.model.chr3.995
Pintab_Pt1G57510
Pintab_Pt1G70920
Pintab_Pt7G23830
Seqgig_SEGI_09451
Wolnob_Wono159310.5
Ginbil_evm.model.chr11.1120
Pintab_Pt5G22840
Seqgig_SEGI_26430
Seqgig_SEGI_20561
Wolnob_Wono424970.5
Wolnob_Wono424990.1
0.2

# h

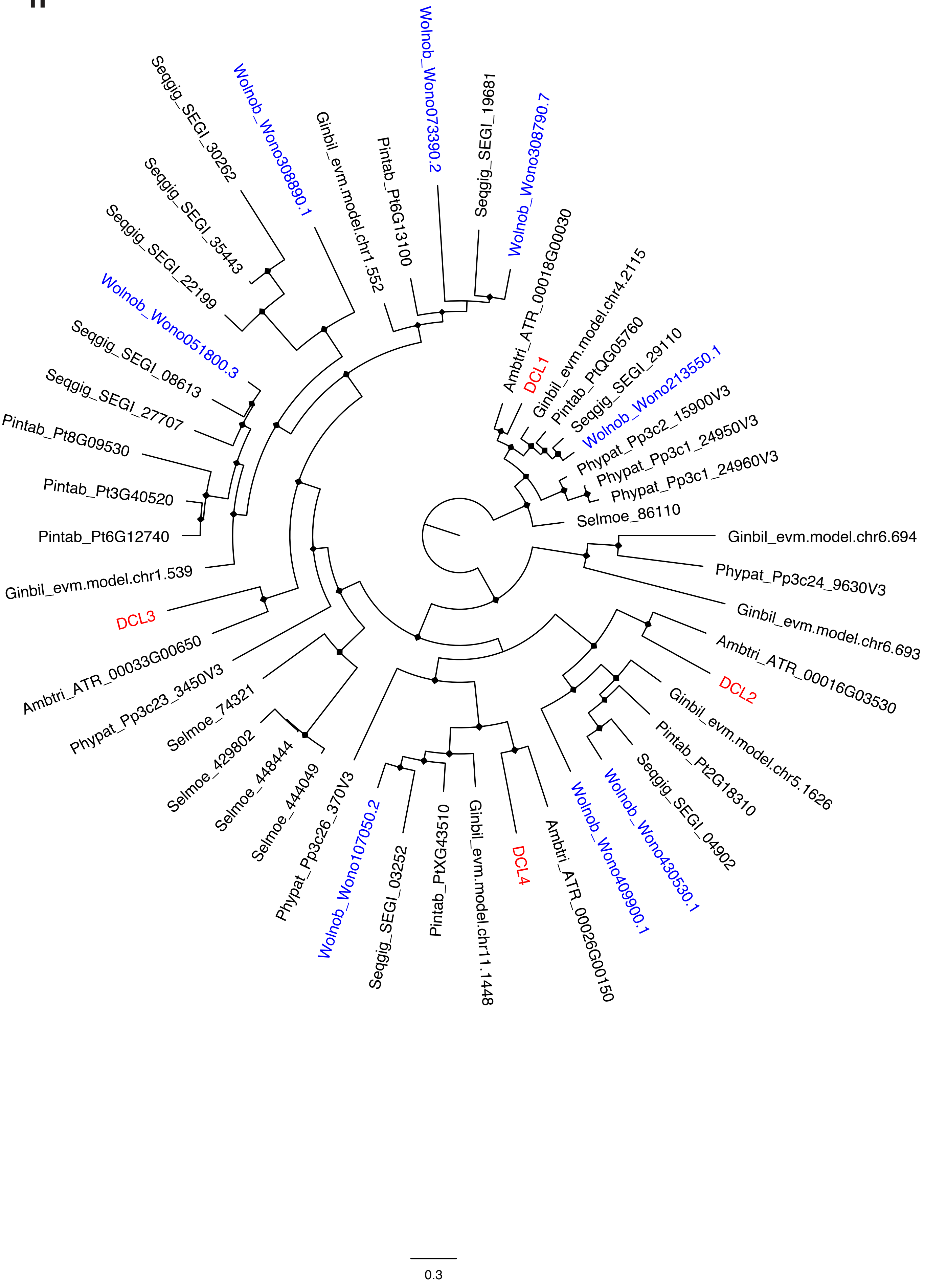

Seqgig_SEGI_30262
Wolnob_Wono308890.1
Ginbil_evm.model.chr1.552
Pintab_Pt6G13100
Wolnob_Wono073390.2
Seqgig_SEGI_19681
Wolnob_Wono308790.7
Ambtri_ATR_00018G00030
DCL1
Ginbil_evm.model.chr4.2115
Pintab_PtQG05760
Seqgig_SEGI_29110
Wolnob_Wono213550.1
Phypat_Pp3c2_15900V3
Phypat_Pp3c1_24950V3
Phypat_Pp3c1_24960V3
Selmoe_86110
Ginbil_evm.model.chr6.694
Phypat_Pp3c24_9630V3
Ginbil_evm.model.chr6.693
Ambtri_ATR_00016G03530
DCL2
Ginbil_evm.model.chr5.1626
Pintab_Pt2G18310
Seqgig_SEGI_04902
Wolnob_Wono430530.1
Wolnob_Wono409900.1
Ambtri_ATR_00026G00150
DCL4
Ginbil_evm.model.chr11.1448
Pintab_PtXG43510
Seqgig_SEGI_03252
Wolnob_Wono107050.2
Phypat_Pp3c26_370V3
Selmoe_444049
Selmoe_448444
Selmoe_429802
Selmoe_74321
Phypat_Pp3c23_3450V3
Ambtri_ATR_00033G00650
DCL3
Ginbil_evm.model.chr1.539
Pintab_Pt6G12740
Pintab_Pt3G40520
Pintab_Pt8G09530
Seqgig_SEGI_27707
Seqgig_SEGI_08613
Wolnob_Wono051800.3
Seqgig_SEGI_22199
Seqgig_SEGI_35443
0.3

i

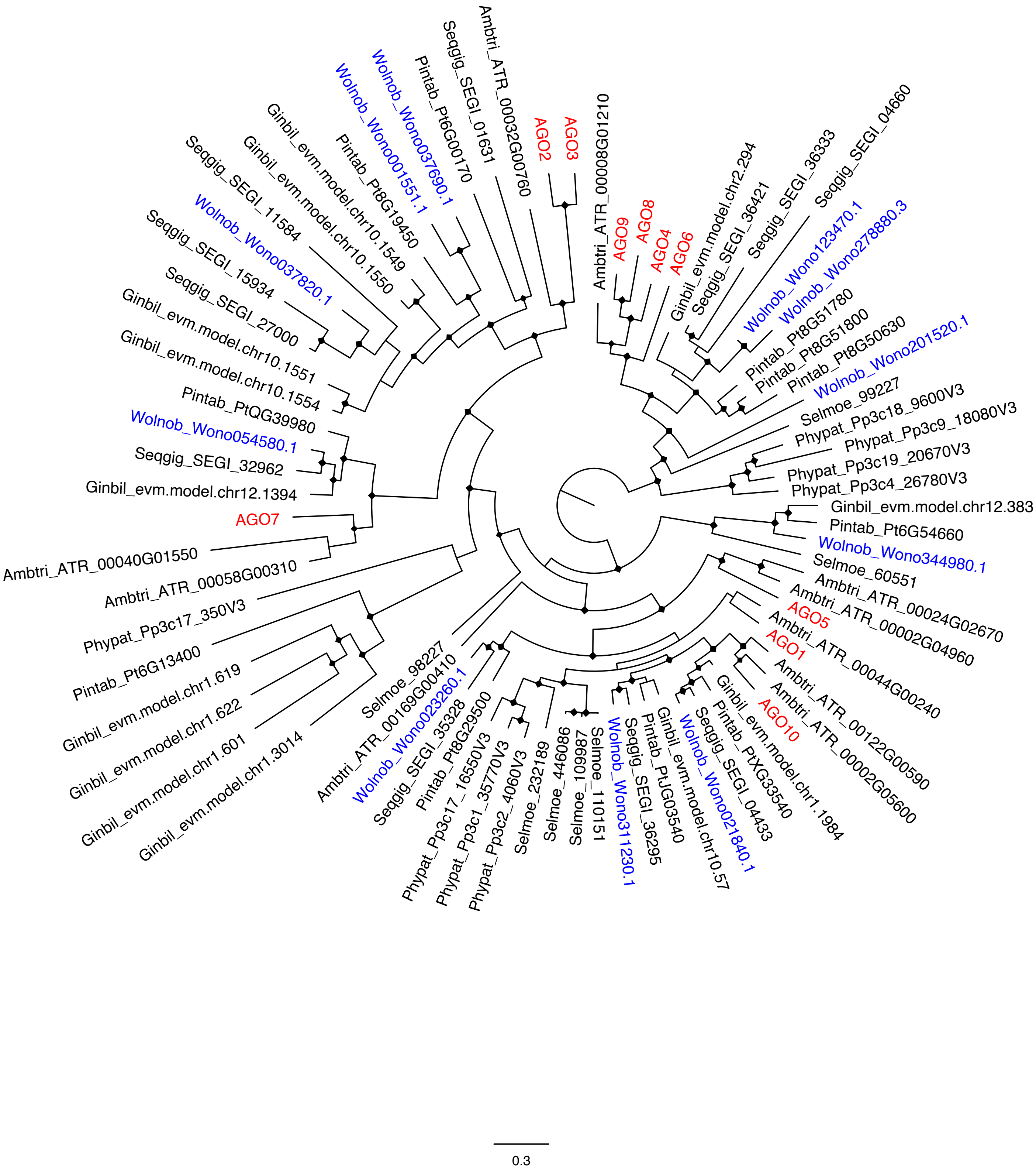
AGO2
AGO3
AGO9
AGO8
AGO4
AGO6
AGO7
AGO5
AGO1
AGO10
Ambtri_ATR_00008G01210
Ambtri_ATR_00032G00760
Seqgig_SEGI_01631
Pintab_Pt6G00170
Wolnob_Wono037690.1
Wolnob_Wono001551.1
Pintab_Pt8G19450
Ginbil_evm.model.chr10.1549
Ginbil_evm.model.chr10.1550
Seqgig_SEGI_11584
Wolnob_Wono037820.1
Seqgig_SEGI_15934
Seqgig_SEGI_27000
Ginbil_evm.model.chr10.1551
Ginbil_evm.model.chr10.1554
Pintab_PtQG39980
Wolnob_Wono054580.1
Seqgig_SEGI_32962
Ginbil_evm.model.chr12.1394
Ambtri_ATR_00040G01550
Ambtri_ATR_00058G00310
Phypat_Pp3c17_350V3
Pintab_Pt6G13400
Ginbil_evm.model.chr1.619
Ginbil_evm.model.chr1.622
Ginbil_evm.model.chr1.601
Ginbil_evm.model.chr1.3014
Selmoe_98227
Ambtri_ATR_00169G00410
Wolnob_Wono023260.1
Seqgig_SEGI_35328
Pintab_Pt8G29500
Phypat_Pp3c17_16550V3
Phypat_Pp3c1_35770V3
Phypat_Pp3c2_4060V3
Selmoe_232189
Selmoe_446086
Selmoe_109987
Selmoe_110151
Wolnob_Wono311230.1
Seqgig_SEGI_36295
Pintab_PtJG03540
Ginbil_evm.model.chr10.57
Wolnob_Wono021840.1
Seqgig_SEGI_04433
Pintab_PtXG33540
Ginbil_evm.model.chr1.1984
Ambtri_ATR_00002G05600
Ambtri_ATR_00122G00590
Ambtri_ATR_00044G00240
Ambtri_ATR_00002G04960
Ambtri_ATR_00024G02670
Selmoe_60551
Wolnob_Wono344980.1
Pintab_Pt6G54660
Ginbil_evm.model.chr12.383
Phypat_Pp3c4_26780V3
Phypat_Pp3c19_20670V3
Phypat_Pp3c9_18080V3
Phypat_Pp3c18_9600V3
Selmoe_99227
Wolnob_Wono201520.1
Pintab_Pt8G50630
Pintab_Pt8G51800
Pintab_Pt8G51780
Wolnob_Wono278880.3
Wolnob_Wono123470.1
Seqgig_SEGI_04660
Seqgig_SEGI_36333
Seqgig_SEGI_36421
Ginbil_evm.model.chr2.294
0.3

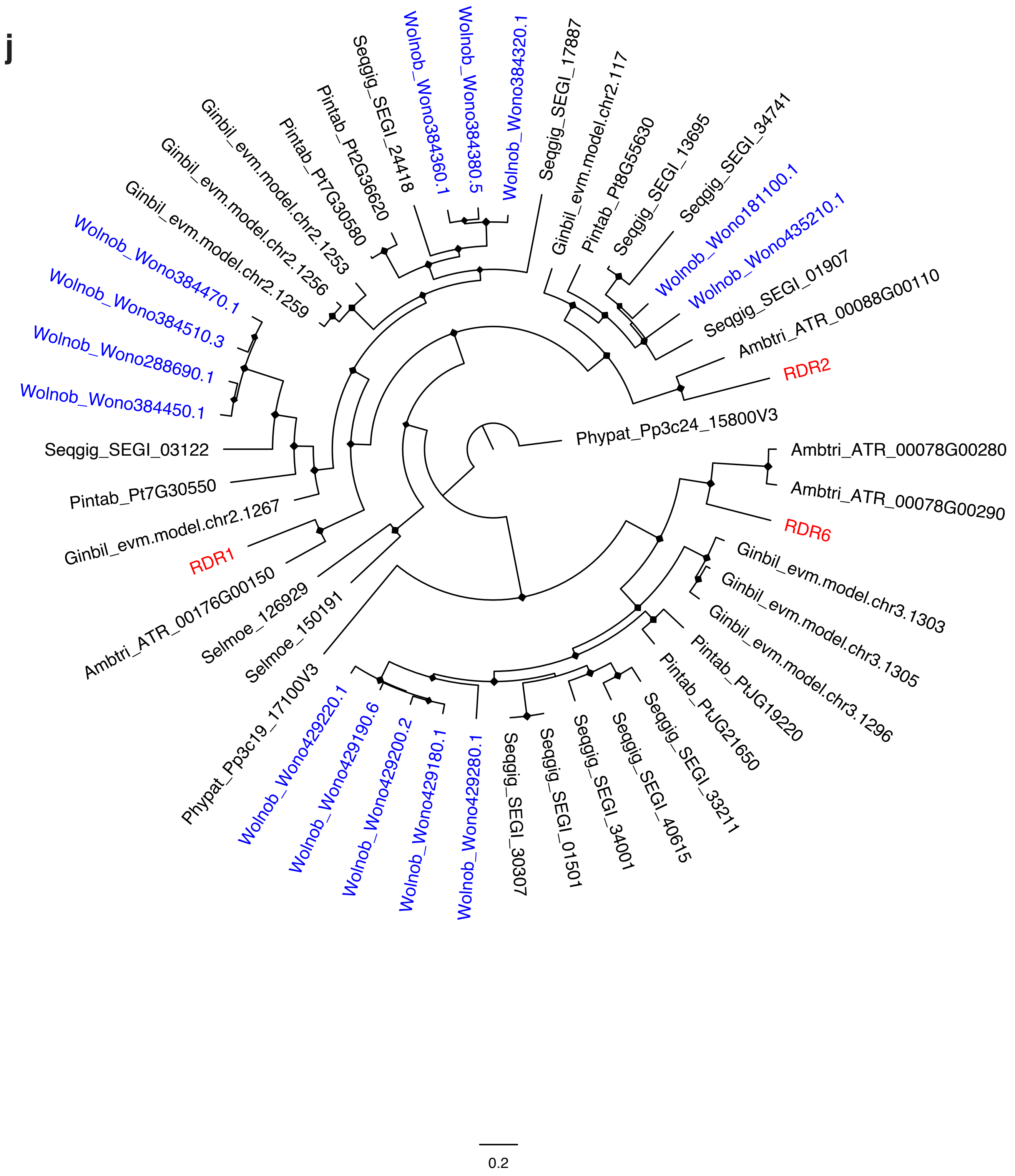


**Supplementary Figure 5 | Phylogenetic reconstruction (maximum likelihood) of select gene families. a,** AP2/ANT across land plants. Sequences of Wollemia nobilis have been found in the euAP2clade and in several ANT clades. **b,** ERF lineage. Several Wollemia nobilis homologs were found in the different clades of this gene lineage. **c,** APETALA2 (AP2) / ETHYLENE RESPONSIVE FACTOR (ERF) large gene family across land plants. Sequences of Wollemia nobilis have been found in all the major clades. In blue are the gene homologs belonging to the AP2/AINTEGUMENTA (ANT), and in black are RAV and ERF gene homologs. **d,** NLR gene family; orthologs are color-coded by predicted subfamily based on a homology search against the RefPlantNLR database (https://doi.org/10.1371/journal.pbio.3001124). The largest subfamily is the TIR-NLR (Toll/Interleukin-1 receptor) subfamily (green) which includes ~63% of the family members. The RWP8 domain-containing CCR subfamily includes 36 members (purple). The CC domain-containing subfamily includes 55 members which are further sub-classified into the CC (blue) and the CCG10 lineage (yellow) with 40 and 15 members, respectively. **e,** Defensin gene family. **f,** CMT gene family. Arabidopsis thaliana orthologs are labeled in red with their common acronyms. Wollemia nobilis orthologs are labeled in blue. Diamonds indicate branches with high bootstrap support (≥80%). **g,** DRM gene family. **h,** DCL gene family. **i,** AGO gene family. **j,** RDR gene family.

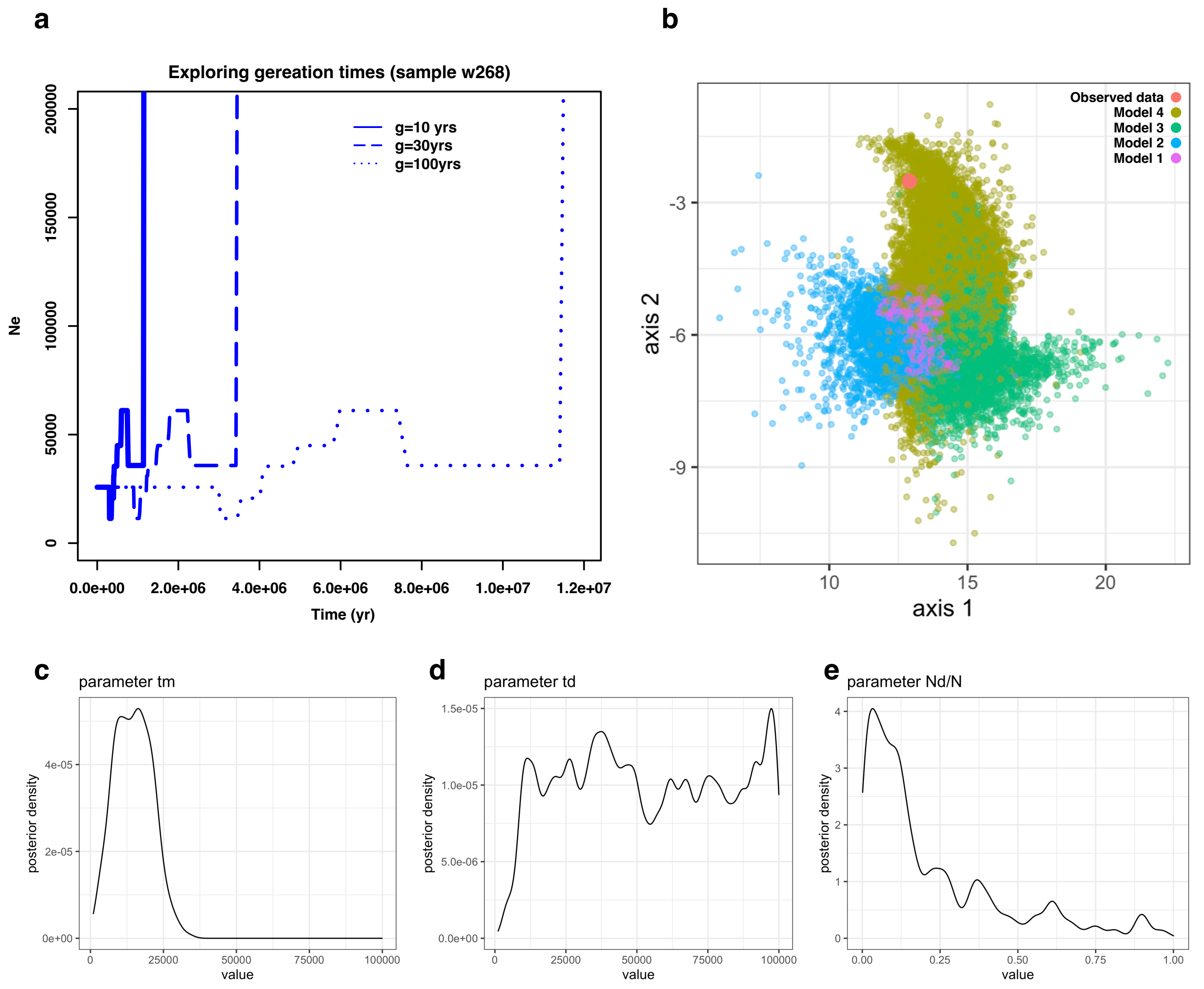


**Supplementary Figure 7 | Demographic history of the Wollemi pine based on whole genomes and DArTSeq data. a,** Demographic history based on a whole genome of the Wollemi pine (sample NSW1032268), using three different generation times. **b,** Principal component analysis of pseudo-observed data simulated under four demographic scenarios, based on DArTSeq data of two Wollemi pine populations. The beige, green, blue and purple dots correspond to models 1 through 4, respectively (SuppFig 9). The observed data is in red. **c-e,** Posterior distribution of parameter estimates from the best model for two Wollemi pine populations. **c,** Time (in generations) of split between two extant populations. **d,** Time of population size decreases in the ancestral population. **e,** Ratio between ancestral population sizes post and pre decrease.

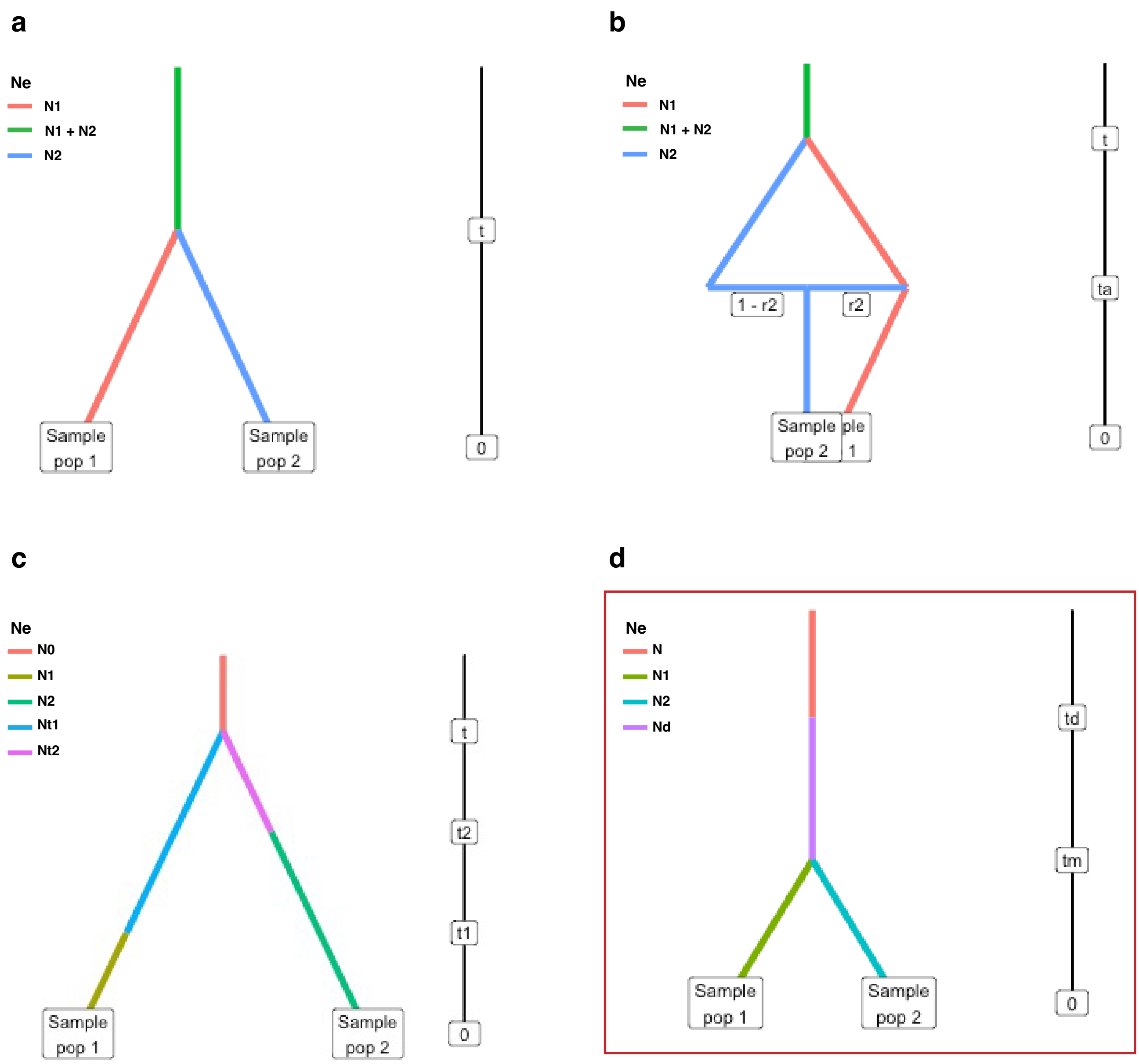


**Supplementary Figure 8 | Diagrams representing four demographic scenarios compared for populations S1 and S2 of the Wollemi pine, with a red box around the best model. a,** simple ancestral population split model with no associated population size change (scenario 4). **b,** ancestral population split followed by admixture between the daughter populations (scenario 3). **c,** split followed by independent population size decreases in the daughter populations (scenario 2). **d,** decrease in the ancestral population followed by split into daughter populations (scenario 1).
